# FROM CANCER MOLECULAR SUBTYPE TO AI HYPE: BENCHMARKING AI IN CANCER MOLECULAR SUBTYPING

**DOI:** 10.1101/2025.03.10.642355

**Authors:** Ahtisham Fazeel Abbasi, Muhammad Sajjad, Muhammad Nabeel Asim, Sebastian Vollmer, Andreas Dengel

## Abstract

**Background:** Cancer molecular subtype classification is an essential component of precision oncology which provides insights into cancer prognosis and guides targeted therapy. Despite the growing applications of AI for cancer molecular subtype classification, challenges persist due to non-standardized dataset configurations, diverse omics modalities, and inconsistent evaluation measures. These issues limit the comparability, reproducibility, and generalizability of AI classifiers across different cancers and hinder the development of robust and accurate AI-driven tools.

**Results:** This study benchmarks 35 unique AI classifiers across 153 datasets, covering 8 omics modalities and 20 different cancers. Particularly, it investigates 6 different research questions, and based on comprehensive performance analyses of the 35 AI classifiers it elucidates the research questions with the following answers: (i) Out of 17 different configurations for 5/8 omics modalities, RPPA (RPPA), Gistic2-all-data-by-genes (CNV), HM27 (Meth), and HiSeqV2-exon (Exon) configurations consistently yield better performance; (ii) In terms of 8 omics modalities, RNASeq, miRNA, CNV, and Exon generally achieve higher macro-accuracy compared to Meth., Array, SNP and RPPA; (iii) SNP and RPPA modalities are prone to biases due to technical noise and data imbalance; (iv) Traditional machine learning (ML) models (SVM, XGB, HGB) perform best on small and low-dimensional datasets, while deep learning (DL) models (ResNet18, CNN, NN, MLP) excel on large and high-dimensional datasets; (v) SVM achieves the highest mean macro-accuracy across all classifiers, with NN, ResNet18, DEEPGENE, and MLP also demonstrate strong performance; and (vi) DL classifiers show superior macro accuracy as compared to ML classifiers in 12 out of 20 cancers.

**Conclusions:** The findings offer key insights to guide the development of standardized, robust, and efficient AI-driven pipelines for cancer molecular subtype classification. This study enhances reproducibility and facilitates better comparison across AI methods, ultimately advancing precision oncology.

**Key Points:**

- This study benchmarks 35 unique AI classifiers, ranging from simpler ML models such as Support Vector Machines (SVM), Histogram-Based Gradient Boosting (HGB), and K-Nearest Neighbors (KNN), to complex DL classifiers including Convolutional Neural Networks (CNNs), computer vision models like DenseNet and ResNet, sequential models such as Recurrent Neural Networks (RNN), Gated Recurrent Units (GRU), Long Short-Term Memory networks (LSTM), and their hybrid combinations (e.g., CNN-LSTM, CNN-GRU), as well as transformer-based models, across 153 datasets spanning 8 omics modalities and 20 cancers. It identifies optimal data configurations and evaluates the performance of these classifiers in cancer molecular subtype classification.
- The study highlights biases in specific omics modalities: SNP, RPPA, and Array exhibit higher variability and precision-recall imbalances, while RNASeq, miRNA, Exon, and CNV deliver more consistent and reliable results.
- ML models (e.g., SVM, XGB, HGB) demonstrate strong performance on smaller datasets with fewer features, whereas DL models (e.g., ResNet18, CNN, NN, MLP, and DEEPGENE transformer) excel in handling high-dimensional datasets with large sample sizes.
- The findings provide critical insights for developing robust, standardized AI pipelines for precision oncology, enhancing reproducibility and enabling meaningful cross-method comparisons.

## 1 Introduction

Cancer is a group of diseases characterized by uncontrolled growth and spread of cells that can invade nearby tissues and other parts of the body (metastasis) [1]. According to the World Health Organization (WHO), more than 200 cancers [2] have been identified with different characteristics, causes, and symptoms [3]. These cancers are categorized based on their origin, structure, and molecular characteristics into several major groups: Carcinomas, Sarcomas, Leukemias, Lymphomas, Neuroendocrine Tumors, and mixed Tumors that vary significantly in terms of origin and mechanism of action [4].

Cancers claim almost 10 million lives each year, and around 19.3 million new cancer cases are diagnosed, making it the second most common cause of death worldwide [5]. These high mortality rates and diagnostic challenges stem from its complex nature and molecular diversity since each cancer presents unique characteristics that complicate diagnosis and treatment [6]. For instance, as multiple cancers are asymptomatic in their early stages, symptoms often mimic other common conditions, and each cancer can have various molecular subtypes requiring distinct treatment approaches. Moreover, the same cancer type can behave differently in individual patients [7] which highlights the critical need to go beyond cancer detection and address the complexities of understanding molecular subtypes and behaviors.

The complexity of understanding cancer molecular subtypes led to Cancer Genome Atlas (TCGA) [8], a landmark project that revolutionized the understanding of cancer’s molecular basis [9]. TCGA has comprehensively mapped the genomic changes in major cancer types, revealing distinct molecular subtypes within what were previously thought to be single diseases. For example: Breast cancer has been classified into at least four major molecular subtypes (Luminal A, Luminal B, HER2-enriched, and Basal-like) Glioblastoma has been categorized into Classical, Mesenchymal, Neural, and Proneural subtypes. These molecular subtypes explain why patients with seemingly similar cancers can have vastly different outcomes and treatment responses.

Traditionally, cancer molecular subtypes are identified through a combination of wet lab experiments and computational analyses. Wet lab experiments involve techniques like genomic and RNA sequencing, epigenomic profiling, and proteomics to generate raw biological data from patient samples [10]. These methods provide valuable insights into genetic mutations, gene expression patterns, regulatory modifications, and protein levels, which are essential for understanding the molecular characteristics of different cancer molecular subtypes. In addition, computational methods such as clustering algorithms, and multi-omics integration are used to identify cancer molecular subtypes and correlate them with clinical outcomes [11]. Although both approaches are valuable for exploring cancer molecular subtypes, wet lab experiments are costly, time-consuming, and prone to variability, while traditional computational tools face challenges with scalability, noise sensitivity, and limited multi-omics integration.

Following the limitations of traditional methods and success of Artificial Intelligence (AI) in various application areas, i.e., omics [12], genomics [13, 14, 15, 16], and proteomics [16, 17, 18], the development of robust AI-based approaches for cancer molecular subtype prediction is an active area of research. Over the last two years, more than 60 AI-based approaches have been developed [19, 20, 21] to predict cancer molecular subtypes more precisely.

Among 60 studies, 15 studies have primarily focused on breast cancer molecular subtypes i.e. Basal-like, Luminal A, Luminal B, and HER2-enriched [22, 23, 24]. This prominence is driven by breast cancer’s high prevalence and the public availability of diverse, and well-curated datasets. Lung cancer ranks second, with 10 studies on subtypes like adenocarcinoma, squamous cell carcinoma, and large cell carcinoma [25, 26, 27]. The remaining studies are distributed across various cancers, including glioblastoma, colorectal cancer, gastric cancer, and leukemia, often tailored around specific subtypes [28, 29, 30, 31].

In terms of datasets, TCGA has been the most frequently utilized resource in over 30 studies for multi-omics data [32, 33, 34, 25]. Other notable sources include the Gene Expression Omnibus (GEO) [35], which has been utilized in 12 studies [36, 37, 38], and cBioPortal in 8 studies [39, 40, 41]. Classical ML classifiers, such as Support Vector Machines (SVM), Random Forest (RF), and Logistic Regression (LR), have been used in approximately 25% of studies [42, 43, 44, 45]. However, DL classifiers have been used in over 50% of studies. Models such as Convolutional Neural Networks (CNNs), and Graph Neural Networks (GNNs) have demonstrated superior performance in handling high-dimensional and multi-modal datasets [22, 46]. In order to deal with the challenges of high dimensionality and noise, over 20 studies have reported the use of methods such as, Principal Component Analysis (PCA), LASSO regression, Variational Autoencoders (VAEs), and t-SNE [47, 48, 49, 50].

Despite significant advancements in AI-based tools for cancer molecular subtype classification, existing studies often focus on individual cancers and offer cancer-specific insights that fail to generalize across diverse cancers. In addition, inconsistent evaluation i.e., varying metrics, and disparate preprocessing strategies, compromise the comparability and reproducibility of results across different studies. Moreover, the heterogeneity in omics modalities and configurations coupled with varied dataset sizes, hinder the development of standardized and robust tools for cancer molecular sub-type classification. Furthermore, to the best of our knowledge, there is no existing study that thoroughly evaluates the predictive performance of AI classifiers for cancer molecular subtype classification by considering the challenges of generalizability across diverse cancer types, standardization of evaluation metrics, and the use of omics data with heterogeneous configurations and varying dataset sizes.

In light of these challenges, there is a pressing need for a comprehensive benchmarking framework that covers multiple cancer types, omics modalities and configurations, and diverse AI classifiers. To address this gap, we present a large-scale benchmarking study that evaluates the predictive performance of 35 ML and DL classifiers, across 153 datasets spanning 8 distinct omics modalities (miRNA, RNASeq, Exon, CNV, SNP, RPPA, and methylation) in terms of 20 different cancers. The objective of this benchmark is to delve into diverse aspects of the cancer molecular sub-type classification, extract and furnish useful insights from diverse experiments with the following different research questions (RQs) and objectives: RQ I) What data configurations are critical for accurate cancer molecular subtype classification across 8 distinct omics modalities? RQ II) How consistently do different omics modalities perform in subtype classification across diverse cancers? RQ III) Are there specific omics modalities that lead to biased predictions, and what factors contribute to this bias? RQ IV) What ML and DL classifiers demonstrate reliable performance across all omics modalities? V) Which specific ML and DL classifiers provide consistent predictive performance with respect to cancers? RQ VI) Among top-performing classifiers in specific cancers, how do ML and DL methods compare in terms of consistency and suitability for cancer molecular subtype classification? We believe answers to these questions will offer critical insights for the research community in identifying the optimal combination of omics modalities, data configurations, and AI classifiers for the development of standardized, robust, and efficient end-to-end predictive pipelines for cancer molecular subtype classification.

## 2 Materials and Methods

This section briefly demonstrates the details of benchmark datasets, ML and DL classifiers, and evaluation measures. A high level graphical illustration of all three core components is given in Figure 1 and briefly described in the following subsection.

**Figure 1:**
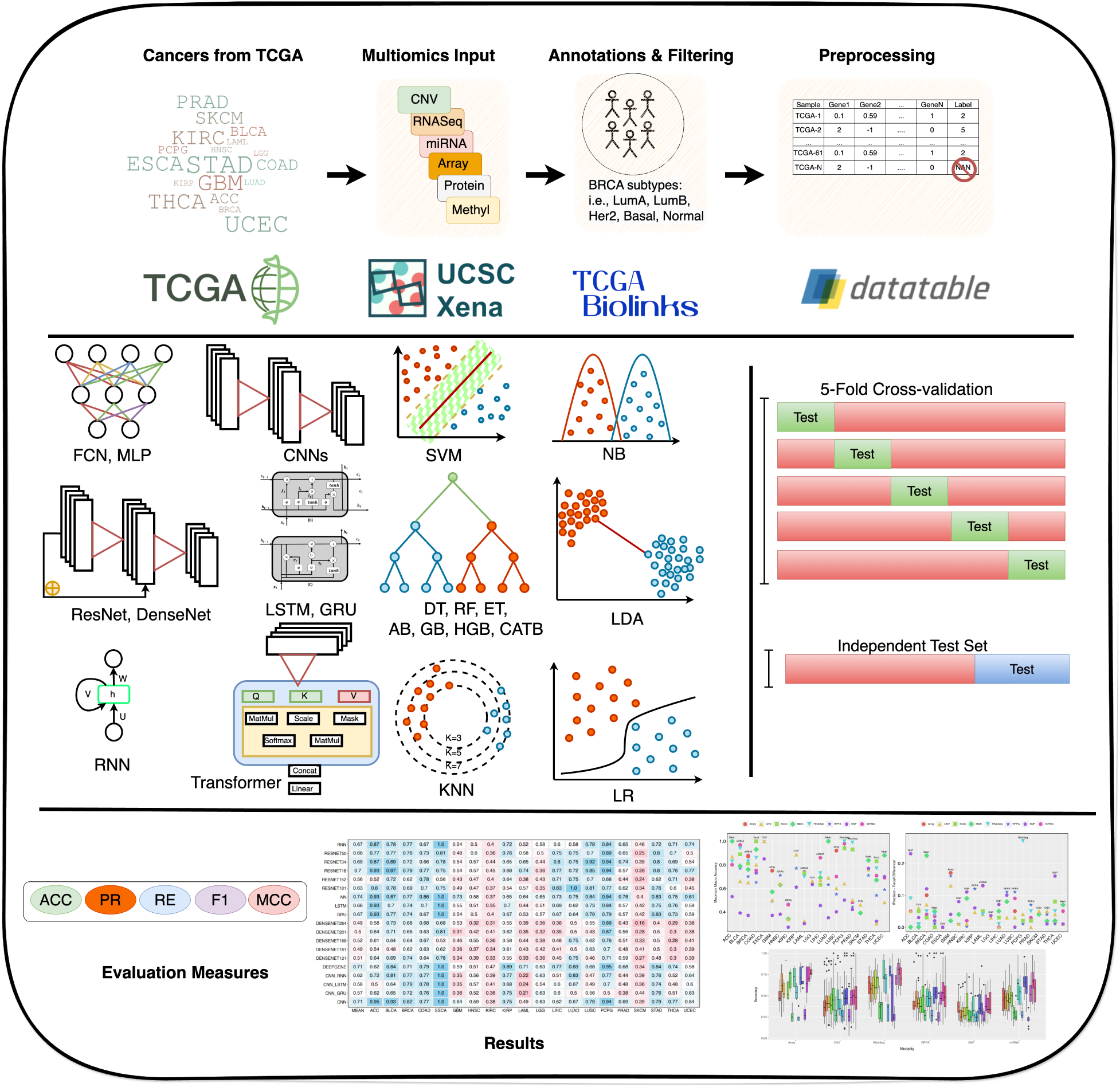
Graphical illustration of the cancer molecular subtype classification benchmarking framework, showcasing dataset preparation, AI classifiers, data splitting strategies, and evaluation methods. In the dataset preparation stage, omics data from TCGA are annotated, filtered, and preprocessed using tools such as UCSC Xena and TCGAbiolinks, covering omics modalities (CNV, RNASeq, miRNA, SNP, RPPA, and methylation). Next, a diverse set of AI classifiers, including 16 ML (e.g., SVM, RF, DT, and AB) and 18 DL (e.g., CNNs, ResNets, DenseNets, LSTMs, and transformers), classifiers are employed. Data splitting is performed using 5-fold cross-validation and independent test sets to ensure robust evaluation. Key performance metrics, including accuracy (ACC), precision (PR), recall (RC), F1-score, Matthews correlation coefficient (MCC), with macro-averaging are applied for comparative analysis across all datasets.

### 2.1 Summary of Cancer Molecular Subtype Classification Benchmark Framework

In recent years, the domain of precision oncology has seen a surge of omics-related datasets for cancer molecular subtype classification. These datasets span multiple omics modalities and are carefully curated to address challenges such as noise, and heterogeneity in molecular profiles. A detailed overview of similar datasets used in this study, spanning 8 distinct modalities and multiple configurations, is provided in section 2.2.

For classification tasks, within the different fields including natural language processing (NLP), computer vision (CV), and genomics, proteomics sequence analysis, ML has witnessed the development of 15 different types of classifiers, namely Naive Bayes (NB) (competent, Bernoulli), Decision Tree (DT) [51], RF [52, 53], Gradient Boosting (GB) [54], Extreme Gradient Boosting (xGB), Histogram-Based Gradient Boosting (HGB) [55], Adaptive Boosting (AB) [56], Light Gradient Boosting Machine (LGBM) [57], Categorical Boosting (CB) [58], K-Nearest Neighbors (KNN) [59], SVM [60], Quadratic Discriminant classifier (QDA) [61], and Logistic Regresison (LR) [62, 63]. Similarly, the DL field has witnessed multifarious architectures built upon core models such as CNNs [64, 65], Recurrent Neural Networks (RNNs) [66], and their combinations, i.e., hybrid models. In addition, transformer-based architectures have emerged as promising architectures based on attention mechanisms to capture long-range dependencies and context in data. The primary objective of this study is to explore the potential of the aforementioned AI classifiers for cancer molecular subtype classification. To achieve this, we conducted in-depth experiments using various ML and DL classifiers. A detailed description of the ML and DL classifiers utilized in this study is provided in the following subsection 2.3.

In order to evaluate the performance of AI-classifiers in the multi-class classification paradigm, 5 evaluation measures are commonly utilized. These evaluation measures encompass accuracy (ACC), precision (PR), recall (RC), F1-score (F1), and Matthews correlation coefficient (MCC) [67]. In our study, AI classifiers are evaluated using these 5 evaluation measures which are briefly discussed in section 2.4.

### 2.2 Datasets

In the pursuit of establishing a robust foundation for the benchmark study of cancer molecular subtype classification, the selection of appropriate datasets plays a pivotal role. The choice of datasets directly impacts the accuracy, robustness, and generalizability of AI classifiers. Poorly selected datasets can result in inappropriate comparisons, biased models, misleading insights, and unreliable decisions in clinical or research settings. Therefore, it is imperative to select datasets that are both well-annotated and representative of the complex nature of cancer molecular subtypes.

Among publicly available resources for cancer molecular subtypes, TCGA offers a comprehensive collection of multiomics datasets that encompasses a broad spectrum of cancer types and subtypes. It is chosen for two key reasons: first, it provides extensive diversity and coverage of cancers and subtypes, which are essential for training and evaluating AI classifiers; second, the datasets are publicly available, well-annotated, and rigorously curated, making them an ideal resource for benchmarking ML and DL classifiers [68, 69, 70].

Table 1 shows 153 benchmark datasets spanning 8 omics modalities and 17 unique configurations, in terms of 20 cancers which are collected from TCGA using UCSC Xena browser. The omics modalities include copy number variation (CNV), DNA methylation, gene expression (RNASeq), microRNA expression (miRNA), single-nucleotide polymorphisms (SNPs), DNA methylation (Meth.), Exon expression (Exon), and protein expression (RPPA). Each modality captures distinct biological features, contributing to the comprehensive characterization of cancer molecular subtypes. A detailed description of these modalities, corresponding configurations, and their technical characteristics are summarized in Table 2. In order to ensure accurate mapping of samples to labels, each dataset is processed using TCGABiolinks to map samples to their corresponding cancer subtype [71]. Datasets with fewer than 70 samples are excluded to ensure sufficient sample size for meaningful analyses and reliable model evaluation.

**Table 1:**
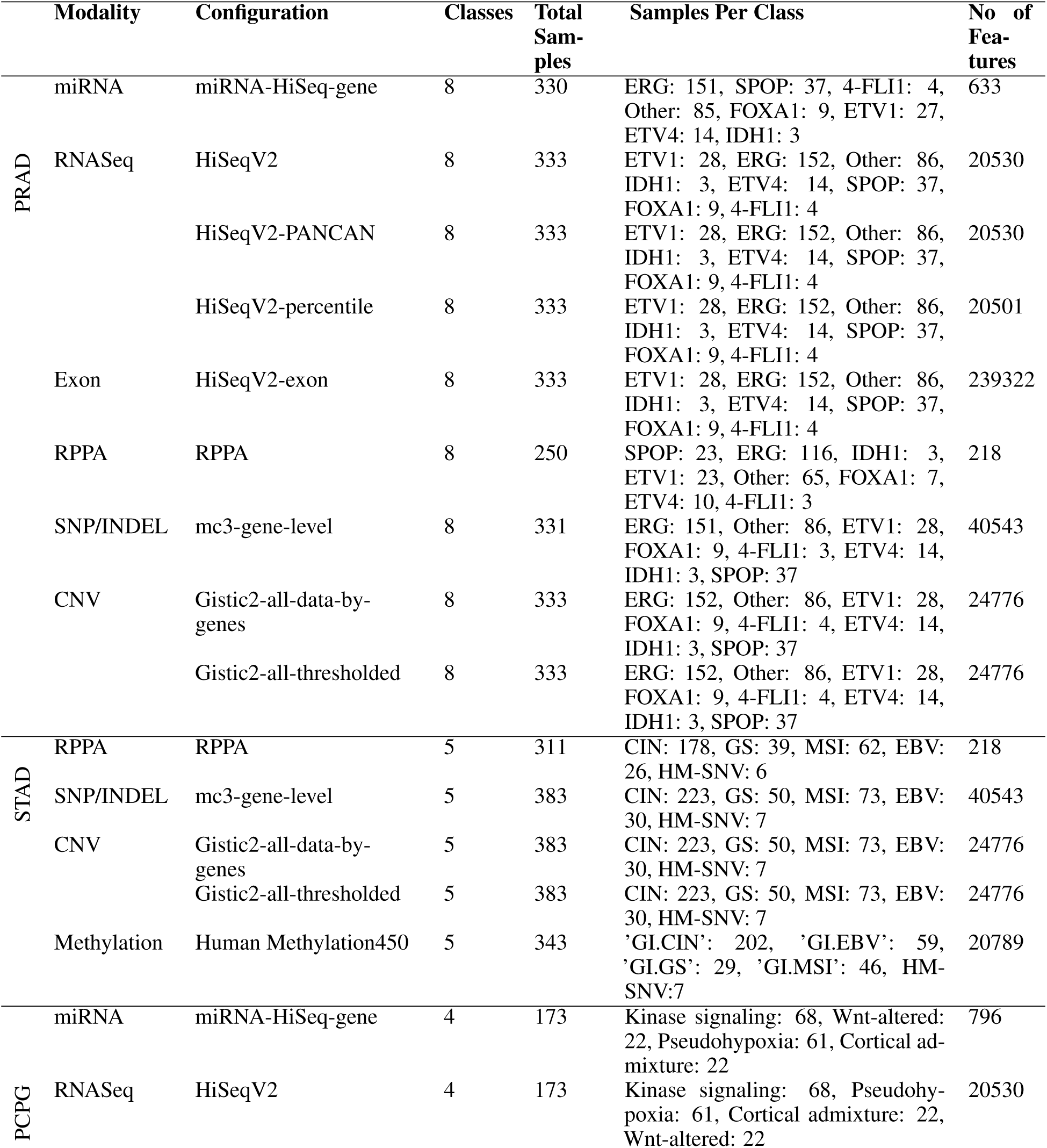
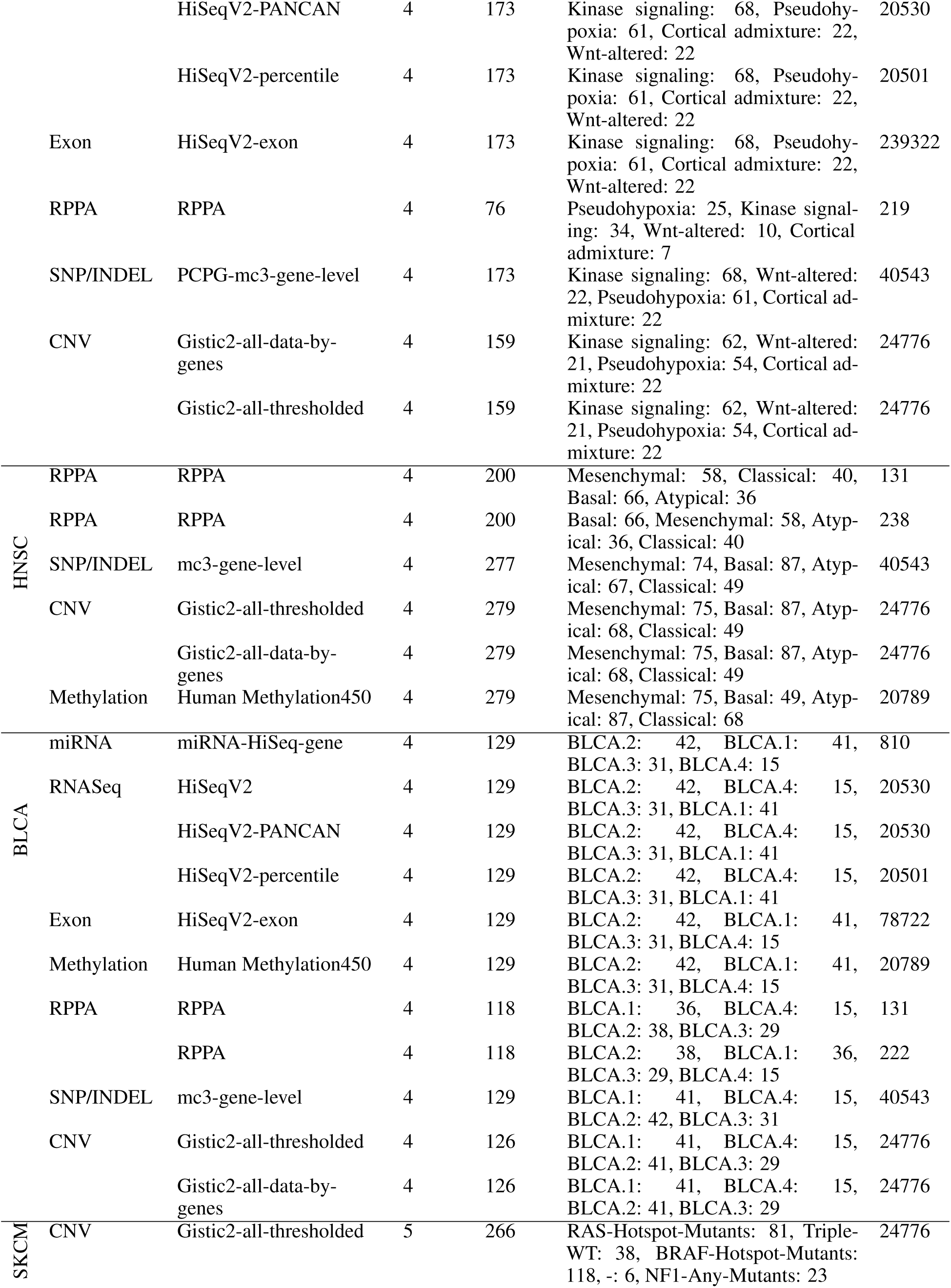
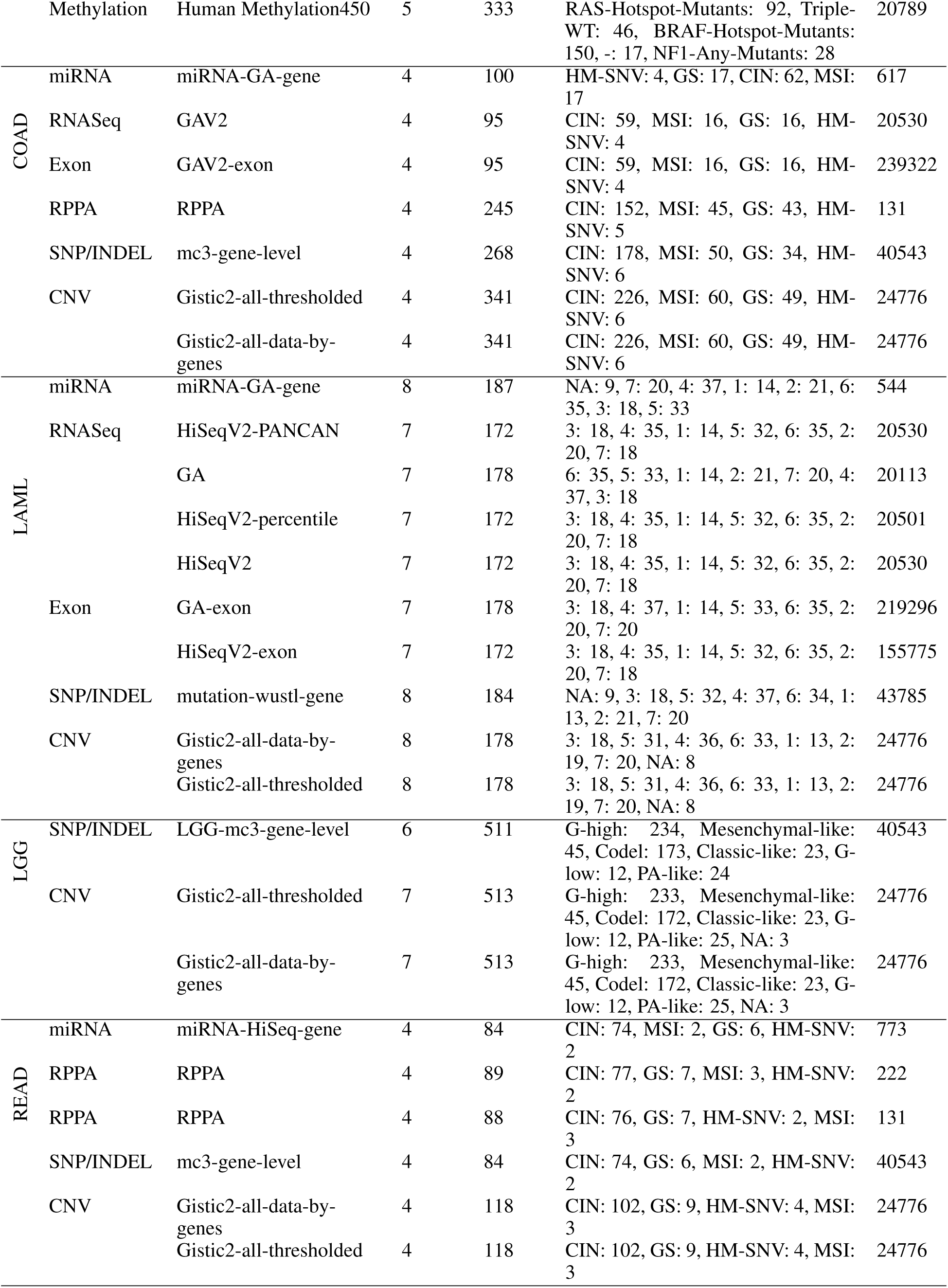
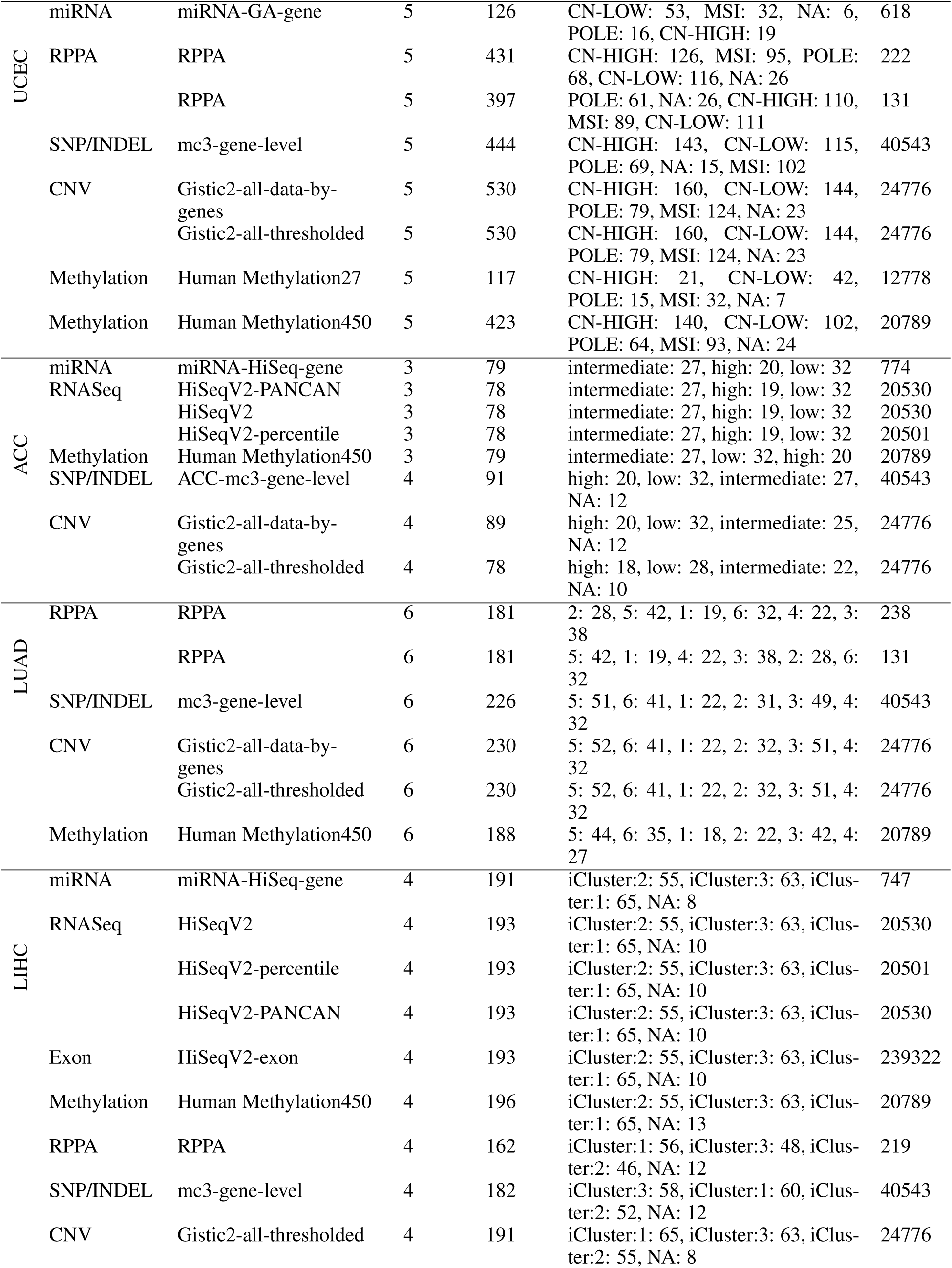
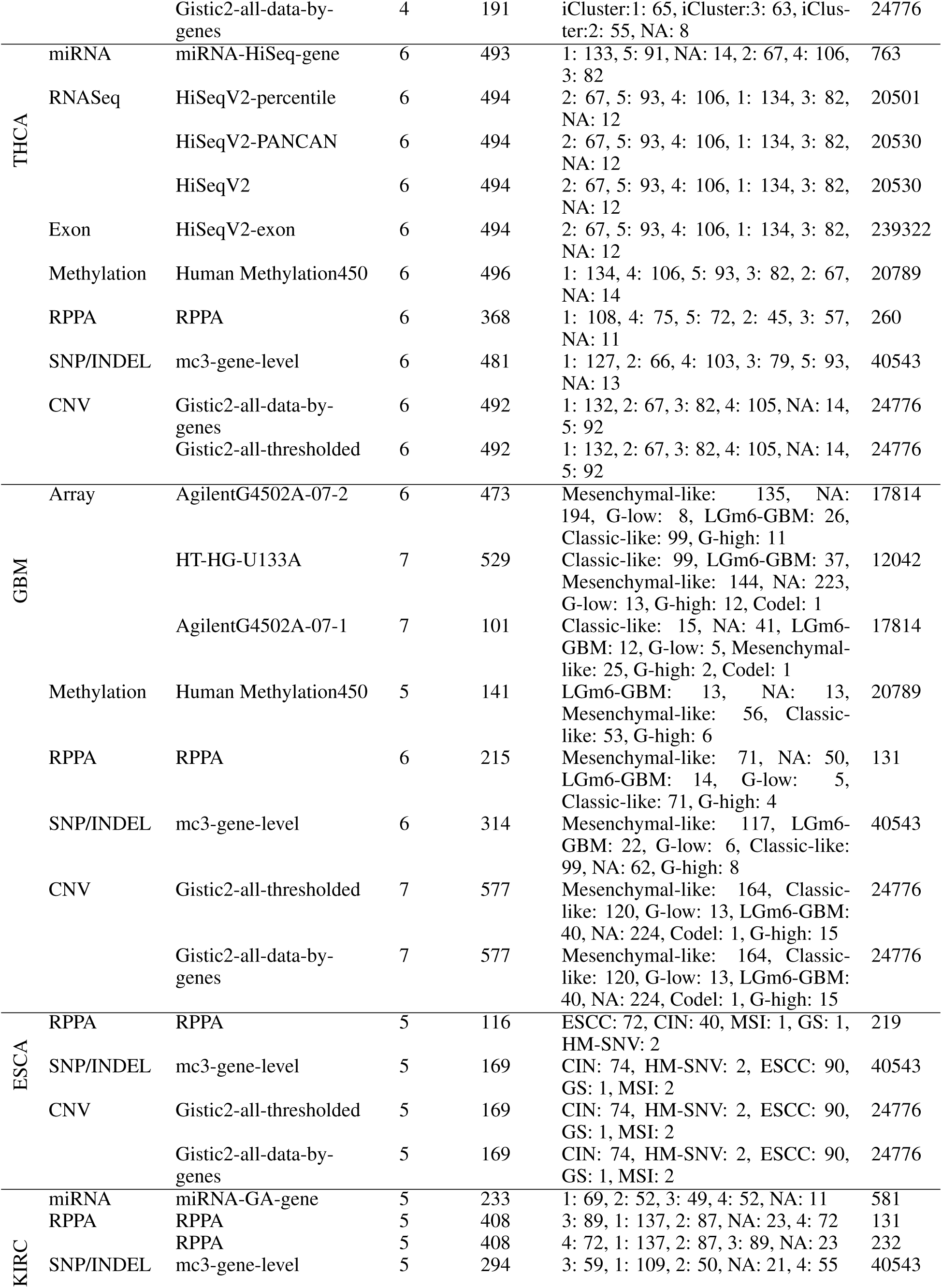
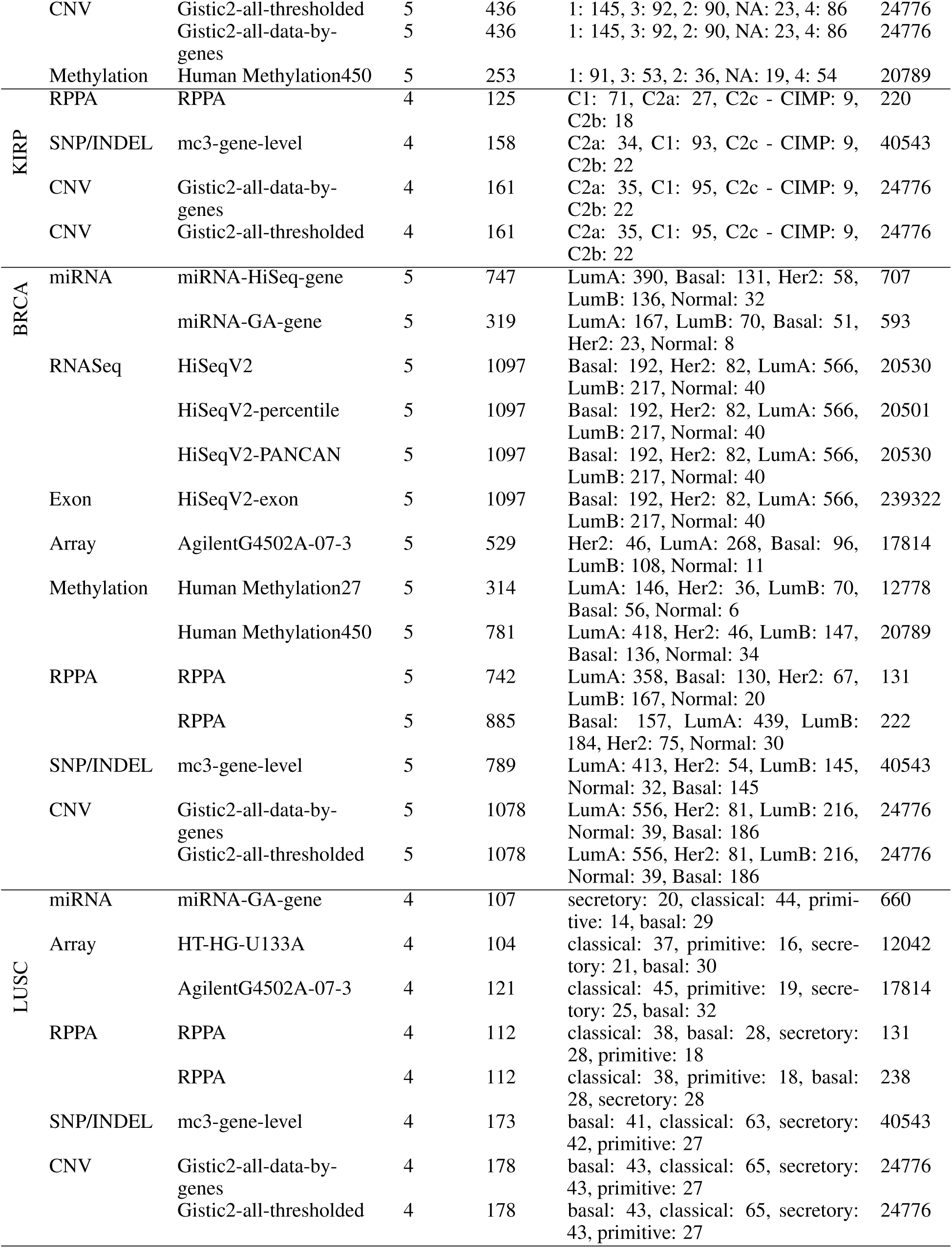
A comprehensive collection of 155 benchmark datasets for cancer molecular subtype classification that encompasses 20 distinct cancers and spans 17 unique data configurations across 8 omics modalities.

**Table 2:**
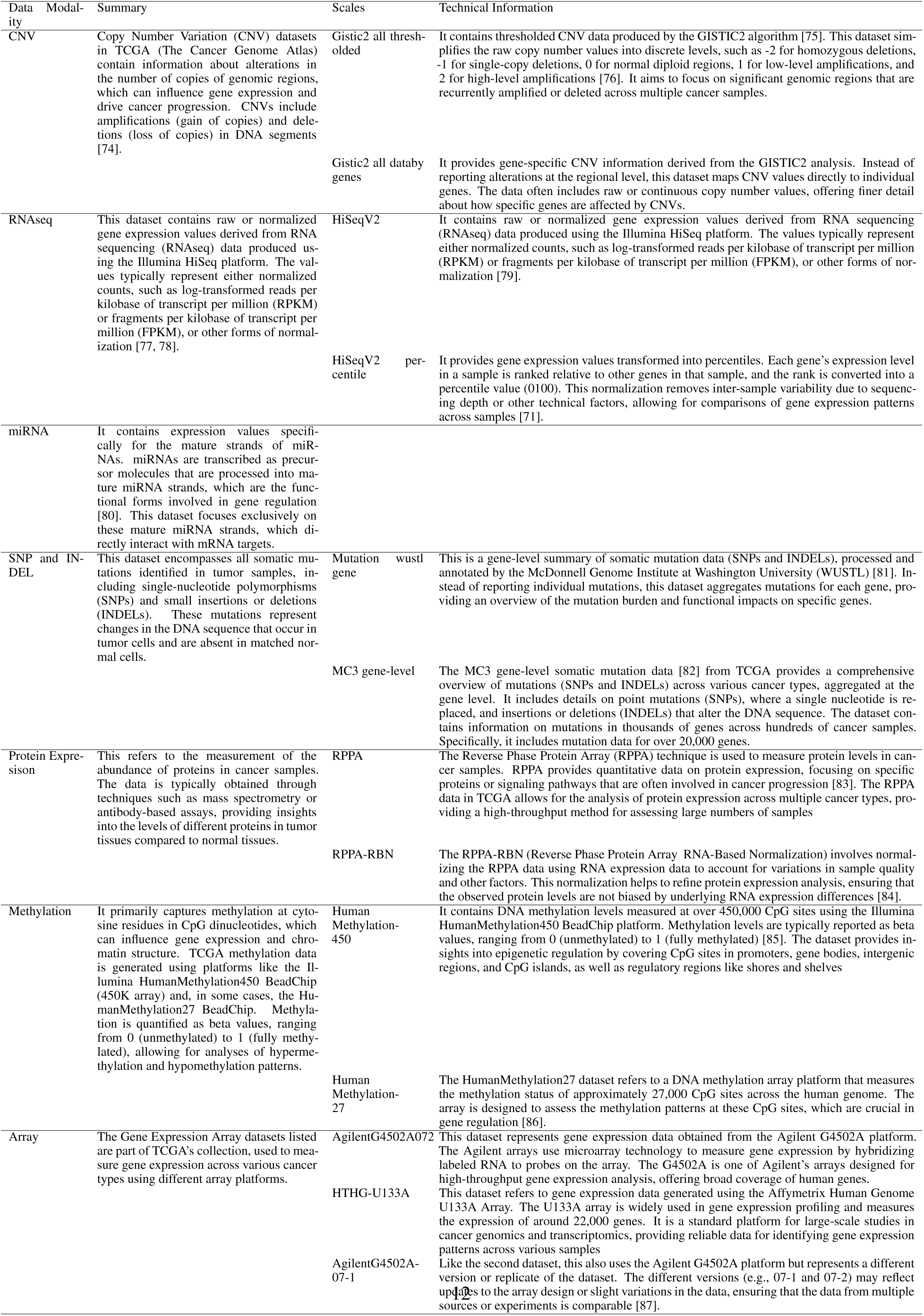
Description of diverse configurations of 8 distinct omics modalities.

The selected datasets encompass a diverse range of sample sizes and subtype distributions which reflect the inherent variability in cancer molecular subtype classification. On average, each dataset includes approximately 328 samples, with BRCA being the most samples i.e., 1,097, while ACC represents the lower end of the spectrum with only 91 samples. This disparity underscores the challenge of modeling rare cancers effectively. In addition, there is a subtype imbalance across all datasets. For instance, in BRCA, the Luminal A subtype has 566 samples, whereas the Normal subtype is significantly underrepresented with just 40 samples. Similarly, in PRAD, the ERG subtype encompasses 152 samples, while FOXA1 accounts for only 9. In addition to differences in sample sizes and subtype distributions, the datasets exhibit considerable variability in features across modalities, which impacts model design and computational requirements. Meth. datasets are the most feature-rich, containing an average of 321,770 features, followed by SNP/INDEL datasets with 40,543 features and RNASeq datasets with 19,552 features. CNV datasets have a moderate feature count of 24,776 on average, while RPPA datasets are notably compact, averaging just 159 features. Given the extensive feature space of methylation datasets, these datasets undergo preprocessing before their use in classification. The details of the preprocessing steps are provided in Supplementary File 1.

This diverse and curated dataset collection provides a solid benchmark for evaluating AI classifiers in cancer molecular subtype classification. It enables a comprehensive assessment of how well AI approaches address the challenges posed by data heterogeneity, subtype imbalance, and modality-specific complexity.

### 2.3 Classifiers

To benchmark the performance of AI classifiers for cancer molecular subtype classification across 153 different benchmark datasets, we have utilized 35 distinct ML (15) and DL (20) classifiers. These classifiers encompass tree and distance-based, and advanced neural network classifiers.

Naive Bayes (NB) is a probabilistic model that uses Bayes’ theorem to assign class labels based on the conditional probabilities of features given the class label. It assumes independence among features and selects the class with the highest posterior probability [72, 73].

Tree-based models include DT [51], RF [52, 53], and ensemble-based methods like GB [54], HGB [55], and AB [56]. The core working principle of tree-based models revolves around recursively splitting the dataset into subsets based on feature thresholds that maximize a splitting criterion, such as information gain or Gini impurity i.e., DT. RF introduces the concept of an ensemble of DTs, where multiple trees are trained on bootstrapped samples, and their predictions are aggregated, typically using majority voting for classification. Boosting methods like GB, HGB, and AB make small modifications to this principle. These algorithms iteratively train weak learners, often decision stumps, on datasets where misclassified samples are given higher weights. HGB optimizes this process by grouping continuous features into histograms for faster computation. In contrast, AB explicitly adjusts sample weights after each iteration, increasing the importance of misclassified samples and reducing the weight of correctly classified ones.

K-Nearest Neighbors (KNN) [59] assigns class labels by identifying the majority class among the k-nearest neighbors of a given data point based on distance metric such as Euclidean distance. SVM [60, 88] find optimal hyperplanes in the feature space to separate classes by maximizing the margin between them. For complex, non-linear boundaries, SVM uses kernel functions to map data to higher-dimensional spaces where the separation is more straightforward.

DL classifiers encompass Multi-Layer Perceptrons (MLPs) [89], CNNs [90], Dense Networks (DenseNets) [91], Residual Networks (ResNets), RNNs [92], LSTMs [93], Gated Recurrent Units (GRUs) [94], and DeepGene transformer [27]. MLPs are feed-forward networks that process input features through multiple layers of interconnected neurons. Each layer applies weighted transformations followed by activation functions, with weights adjusted using backpropagation to minimize prediction error.

CNNs identify patterns by applying convolutional layers that use filters to extract features and pooling layers that reduce dimensionality while retaining critical information. ResNets build upon the CNN architecture by introducing skip connections, which directly connect input and output layers of residual blocks. Such connections enable gradients to bypass one or more layers, alleviating the vanishing gradient problem commonly encountered in deep networks.ResNets facilitate the convergence of more complex architectures, like ResNet18, ResNet34, and ResNet50, thereby improving the capacity to identify nuanced and detailed patterns in datasets. DenseNets take a complementary approach by connecting each layer to every subsequent layer within a dense block. These connections allow features learned by earlier layers to be reused throughout the network and have improved feature propagation. Variants such as DenseNet121, DenseNet161, and DenseNet169 extend the models capacity to learn complex interactions with fewer parameters compared to traditional deep networks.

RNNs are designed to process sequential data by incorporating loops in their architecture, enabling them to maintain a “memory” of previous inputs. However, traditional RNNs struggle with long-term dependencies due to the vanishing gradient problem. Long Short-Term Memory networks (LSTMs) overcome this limitation by introducing gates (input, forget, and output gates) to regulate the flow of information, allowing them to effectively capture long-term dependencies within sequences. GRUs simplify the LSTM architecture by combining the input and forget gates into a single update gate, reducing computational complexity while maintaining the ability to model sequential dependencies effectively. Hybrid models, such as CNN-LSTM and CNN-GRU, combine convolutional layers with recurrent layers, enabling these architectures to learn both spatial and sequential patterns in structured data.

The DeepGene Transformer [27] architecture builds upon the Transformer encoder model, integrating a multi-head self-attention mechanism with 1D convolutional layers. This hybrid design is specifically optimized for high-dimensional gene expression datasets. The model leverages the self-attention mechanism to capture long-range dependencies and intricate patterns across gene features while using 1D convolutional layers to extract local spatial patterns. This combination enables the model to prioritize biologically meaningful features efficiently without requiring prior feature selection and addresses a common limitation in conventional ML approaches.

### 2.4 Evaluation Measures

Following the evaluation criteria of existing cancer classification studies [95, 96, 97], we evaluate 35 AI classifiers with 4 distinct evaluation measures namely, accuracy (ACC), precision (PR), recall (RC), and F1-score (F1). Each measure is computed using a macro-averaging approach to ensure equal weighting for all cancer molecular subtypes irrespective of their sample sizes.

Macro ACC is calculated as the average of individual ACC scores across all cancer molecular subtypes. For a single subtype, ACC is computed as the ratio of correctly predicted samples to the total samples in that subtype. Macro PR is calculated as the average of the ratio of true positives to total predicted positives for each subtype, with single-subtype PR computed as true positives divided by predicted positives. Macro RC is the average of the ratio of true positives to total actual positives, with single-subtype RC calculated as true positives divided by actual positives. Macro F1-score is the harmonic mean of macro-averaged PR and RC, calculated across all subtypes. For individual subtypes, the F1-score is computed as the harmonic mean of the subtype’s PR and RE.

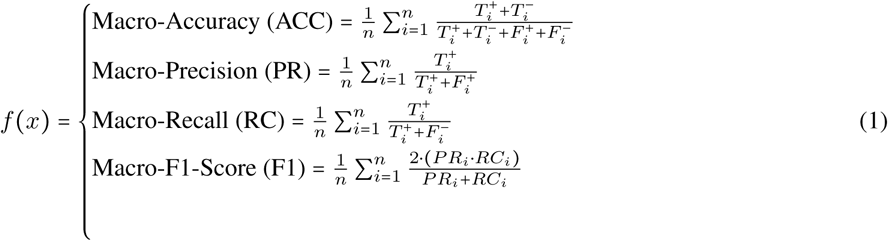

Here, *T_i_*^+^ and *T_i_*^−^ denote the true positive and true negative predictions for subtype *i*, while *F_i_* ^+^ and *F_i_* ^−^ represent false positive and false negative predictions, respectively.

### 2.5 Experimental Setup

To preprocess benchmark datasets, we utilize two different APIs namely, Datatable (https://datatable.readthedocs.io/en/latest/) and TCGABioLinks (https://bioconductor.org/packages/release/bioc/html/TCGAbiolinks.html). Following the evaluation criteria of existing cancer molecular subtype classifiers, we perform experimentation in two different settings namely, 5-fold cross-validation and independent test. In an independent test, 80/20 data-splitting strategy is employed, where 80% of the data is used for training and 20% for testing. Validation data is split from the training data in an 80:20 ratio prior to classifier training. In addition, stratification is applied to ensure that each class is proportionally represented in both the training and test sets [98]. 5-fold cross-validation is also performed with stratification to provide a more comprehensive evaluation of model performance and reduce potential biases introduced by specific data splits [99]. All of the ML classifiers are developed on top of Scikit-Learn v1.3.2 (https://scikit-learn.org/stable). The DL classifiers are developed with Pytorch (https://pytorch.org/). All visualizations are generated using matplotlib v3.8.0 (https://matplotlib.org/) and PlotNine (https://plotnine.org/).

## 3 Results

This section layer by layer unfolds the predictive performance of 35 unique ML and DL classifiers across 153 datasets of 8 distinct omics modalities in terms of 20 different cancers. First, it presents a performance comparison of AI classifiers on 17 unique configurations of 5/8 omics modalities to identify the most effective configurations for each omics modality. Next, it unveils the potential of 8 distinct omics modalities for cancer molecular subtype classification across 20 unique cancers. In addition, it also discusses the performance trends of 35 ML and DL classifiers for 8 omics modalities and 20 different cancers. Finally, it furnishes information about top-performing configuration, modality, and classifier combinations and provides insights into the strengths and weaknesses of ML and DL classifiers for cancer molecular subtype classification. It is important to note that the results presented in this section are derived from independent testing, while 5-fold cross-validation results are presented in Supplementary File 2.

### 3.1 RQ I: What data configurations are critical for accurate cancer molecular subtype classification?

As TCGA provides multiple dataset configurations for different omics modalities as described in section 2.2 and Table 2, selecting the optimal configuration plays a key role in cancer molecular subtype classification. To illustrate this, this subsection answers RQ I by evaluating the classification performance of 17 dataset configurations from 5 omics modalitiesArray, CNV, Meth., RNASeq, and RPPAdue to dataset availability constraints. By assessing ML and DL classifiers on independent test sets across 20 cancers, this study determines the most effective dataset configurations based on macro ACC scores, which offers insights into their suitability for cancer molecular subtype classification.

Figure 2 illustrates the highest macro ACC achieved by ML and DL classifiers across 17 configurations of 5/8 distinct omics modalities, whereas detailed performance scores are provided in Supplementary File 1 Table S1. For CNV, classifiers trained on Gistic2-all-data-genes achieve a higher average macro ACC of 0.624, outperforming Gistic2-all-thresholded, which achieves 0.568. This trend is particularly evident in 13 cancers, including ACC (0.18), BLCA (0.09), BRCA (0.06), GBM (0.1), HNSC (0.13), KIRP (0.147), LAML (0.07), LIHC (0.02), LUAD (0.07), PRAD (0.05), STAD (0.08), THCA (0.03), and UCEC (0.03). In contrast, Gistic2-all-thresholded demonstrates either superior or equivalent performance in 6 cancers: COAD, ESCA, KIRP, LGG (0.01), LUSC (0.01), and PCPG (0.08). Notably, SKCM is excluded from this comparison due to the absence of a Gistic2-all-thresholded dataset. Classifiers trained with Gistic2-all-data-genes achieve better predictive performance due to its granular, continuous representation of gene-level copy number variations, which preserves subtle patterns and rich genomic details critical for cancer molecular subtype classification. Conversely, Gistic2-all-thresholded simplifies data into discrete categories (gain, loss, neutral), effectively reducing noise but potentially discarding critical genomic variability.

**Figure 2:**
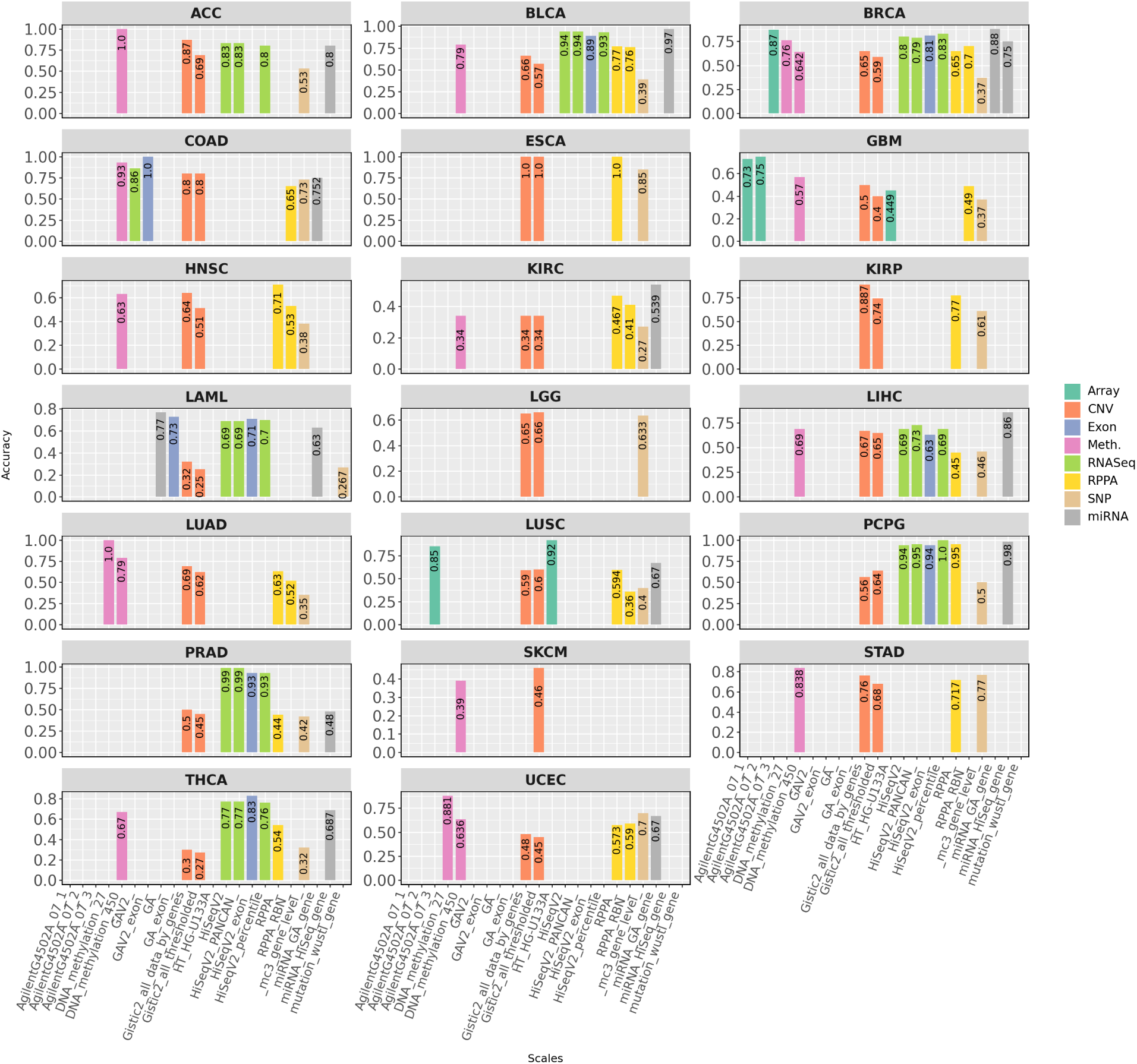
Macro ACC values of top performing classifiers in terms of 20 different configurations of 8 distinct omics modalities across 20 different cancers.

In terms of RNASeq, four configurationsHiSeqV2, HiSeqV2-PANCAN, HiSeqV2-percentile, and GAV2are considered. However, GAV2 is excluded from this analysis due to insufficient data across multiple cancers. A closer examination of macro ACC reveals that classifiers trained on HiSeqV2, HiSeqV2-PANCAN, and HiSeqV2-percentile exhibit nearly identical performance, achieving average macro ACC scores of 0.83, 0.84, and 0.83, respectively, across 8 cancers: ACC, BLCA, BRCA, LAML, LIHC, PCPG, PRAD, and THCA. Notably, COAD, ESCA, GBM, HNSC, KIRC, KIRP, LGG, LUAD, LUSC, SKCM, STAD, and UCEC are excluded from this analysis due to a lack of comparable data. The consistent performance across these RNASeq configurations suggests that they preserve key biological signals while minimizing noise, ensuring robustness in cancer molecular subtype classification regardless of the preprocessing or normalization strategy.

In the context of protein expression, a comparison between RPPA and RPPA-RBN across seven cancers shows that classifiers trained on RPPA achieve a higher average macro ACC (0.627) compared to RPPA-RBN (0.552). Specifically, RPPA outperforms RPPA-RBN in five cancers: BLCA (0.01), HNSC (0.18), LUSC (0.23), KIRC (0.057), and LUAD (0.11). In contrast, RPPA-RBN slightly outperforms RPPA in BRCA (0.05) and UCEC (0.017). The superior performance of RPPA in most cancers may be attributed to its ability to preserve critical protein expression signals without additional transformations. By avoiding normalization, RPPA better captures unique molecular profiles and protein expression variations across different cancers, whereas RPPA-RBN’s normalization process may obscure subtle subtype-specific differences.

For the array modality, 4 configurationsAgilentG4502A-07-1, AgilentG4502A-07-2, AgilentG4502A-07-3, and HTHG-U133Aare evaluated across GBM (AgilentG4502A-07-1 and AgilentG4502A-07-2) and LUSC (AgilentG4502A-07-3 and HTHG-U133A). While this analysis does not encompass all configurations across cancers, it provides valuable insights into their performance in specific cases. For GBM, classifiers trained on AgilentG4502A-07-2 slightly outperform those trained on AgilentG4502A-07-1, with a macro ACC difference of 0.02. This minor improvement may be attributed to more refined data preprocessing in AgilentG4502A-07-2, potentially better aligning with the molecular characteristics of GBM. However, further comparisons across additional cancers and configurations are needed for conclusive insights. For LUSC, classifiers trained on HTHG-U133A outperform those using AgilentG4502A-07-3 by a margin of 0.07. Although this difference is modest, it may be due to HTHG-U133A offering a broader dynamic range or improved feature representation, enhancing the detection of subtype-specific patterns in LUSC. Nonetheless, additional experiments and datasets are required to validate these findings across broader conditions.

For Meth. modality, the analysis is conducted only on BRCA and LUAD, as the two available configurationsHuman-Methylation27 (HM27) and HumanMethylation450 (HM450)are limited to these cancers. In BRCA, classifiers trained on HM27 outperform those using HM450, with a macro ACC margin of 0.118. Similarly, in LUAD, HM27-based classifiers demonstrate superior performance with a macro ACC margin of 0.21. Classifiers trained on HM27 outperform those on HM450 due to a higher signal-to-noise ratio, as HM27 focuses on promoter CpG islands, while HM450 includes many low-variance, sparsely methylated sites that add noise. The lower dimensionality of HM27 (27k vs. 450k features) helps avoid overfitting, whereas HM450s high feature-to-sample ratio and batch effects reduce classifier robustness.

Overall, the analyses of 17 dataset configurations across 5 distinct omics modalities conclude that Gistic2-all-data-genes outperforms Gistic2-all-thresholded for cancer molecular subtype classification in CNV. In RNASeq, HiSeqV2, HiSeqV2-PANCAN, and HiSeqV2-percentile achieve nearly identical performance, which confirms their robustness across cancers. In RPPA, unprocessed data provides better results than RPPA-RBN, as it retains essential protein expression signals. The array modality shows variations based on configuration, while in Meth., HM27 outperforms HM450 due to a higher signal-to-noise ratio and a lower risk of overfitting. These results emphasize the importance of choosing appropriate dataset configurations, as differences in preprocessing and representation methods impact predictive performance of classifiers for cancer molecular subtype classification.

### 3.2 RQ II **&** III) How consistently do different omics modalities perform in cancer subtype classification across diverse cancers?

In the previous subsection, we determined the most effective configurations for each omics modality for cancer molecular subtype classification. Building on that analyses, this subsection evaluates the effectiveness of 8 distinct omics modalities Array, CNV, Meth., RNASeq, RPPA, SNP, Exon, and miRNAin cancer molecular subtype classification across 20 different cancers. To illustrate this, this subsection answers RQ II and III by presenting the maximum macro ACC and minimum PR-RE disparity achieved by an ML or DL classifier for each of the 8 omics modalities. This analysis aims to highlight the strengths and limitations of each omics modality in achieving accurate and robust cancer molecular subtype classification. In addition, the availability of 8 omics modalities is not uniform across all cancers, hence the analyses and conclusions are drawn based on the omics modalities present for each cancer.

Figure 3.a illustrates the maximum macro ACC achieved by the top-performing classifiers for 20 different cancers across 8 omics modalities: Array, CNV, RNASeq, RPPA, SNP, Exon, Meth., and miRNA, with a detailed breakdown of these performance scores provided in Supplementary File 1 Table S2. The performance analysis of omics modalities across 20 different cancers highlights distinct trends i.e., omics modalities with maximum and minimum performances, disparity of PR and RE across omics modalities. Classifiers trained with specific omics modalities achieving maximum performance demonstrate the effectiveness of these modalities for cancer molecular subtype classification, and vice versa. For instance, classifiers trained on miRNA modality perform better in terms of macro ACC across 5/12 cancers (BLCA, BRCA, KIRC, LAML, and LIHC). This trend establishes miRNA as the most informative omics modality overall for cancer molecular subtype classification. Similarly, classifiers trained on Meth. modality achieve maximum performance in 4/13 cancers namely, ACC, LUAD, UCEC, and STAD. Similarly, classifiers trained with CNV modality demonstrate better macro ACC in 4/20 cancers (KIRP, ESCA, SKCM, and KIRP). In addition, classifiers based on Exon modality show better performance in (2/8) cancers namely, COAD, and THCA. Although the RNASeq modality allows classifiers to achieve the highest macro ACC 2/9 cancers (PCPG, and PRAD), these classifiers consistently demonstrate reasonable performance compared to those based on other modalities. Classifiers using Array and RPPA modalities perform best in 3 cancers (Array (2/3): GBM, LUSC; RPPA (1/14): HNSC). Conversely, classifiers trained on SNP modality frequently show the lowest performance, particularly in cancers such as ACC, LAML, RPPA, BLCA, BRCA, KIRP, GBM, KIRC, HNSC, KIRP, LAML, LGG, LUAD, LUSC, PCPG, PRAD, and THCA. This indicates the reduced predictive capacity for RPPA modality in 12 cancers for cancer molecular subtype classification.

**Figure 3:**
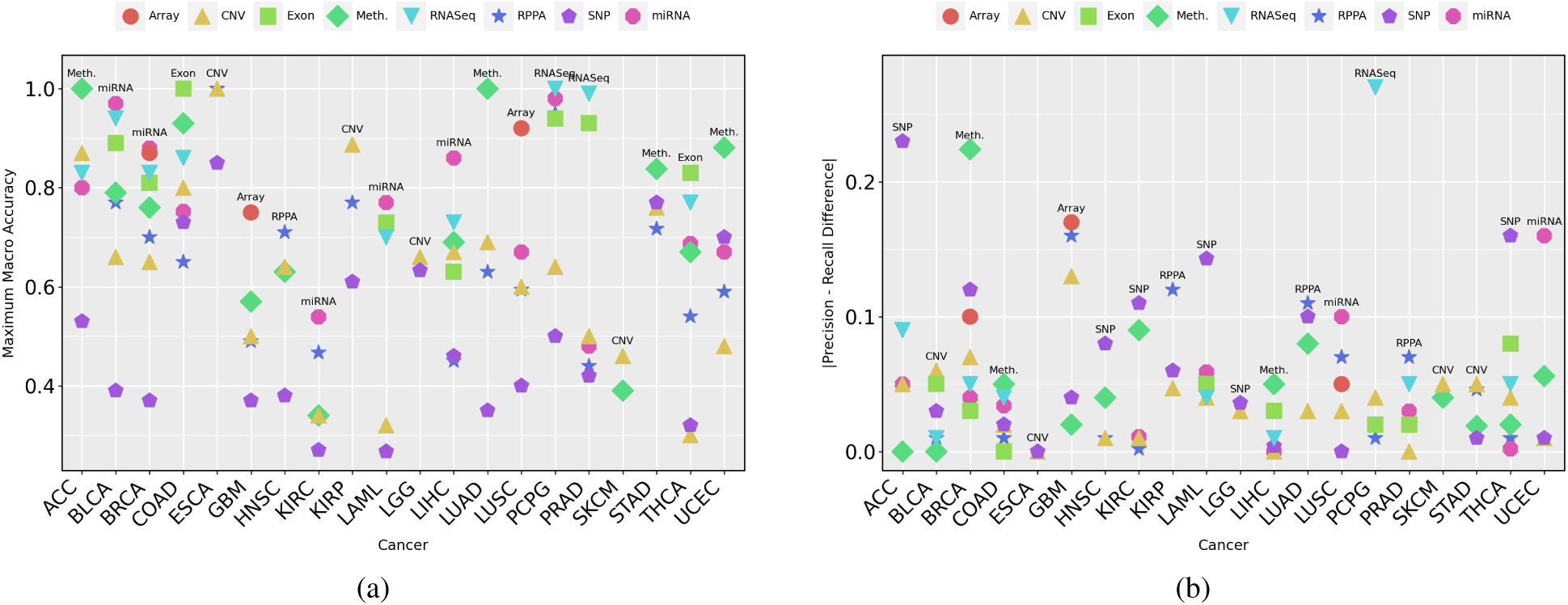
(a) Macro-ACC of top performing classifier-modality combination across 20 different cancers in terms of 7 distinct omics modalities. (b) PR-RC difference of top performing classifier-modality combination for a cancer. A larger PR-RC difference indicates a higher degree of bias in the molecular subtype classification results for the corresponding cancer.

Figure 3.b shows the disparity between macro PR and RE across 20 cancers and 8 omics modalities. This disparity reflects imbalances in classification performance, where PR does not always align with RE which highlights potential biases or limitations in cancer molecular subtype classification. Classifiers using SNP modality exhibit the highest disparity in 5 cancers such as ACC (0.23), KIRC (0.11), LAML (0.14), HNSC (0.08) and TCHA (0.16). This highlights challenges in achieving accurate molecular subtype classification for these cancers, even though SNP-based classifiers may achieve moderate PR in some cases. Similarly, classifiers using RPPA modality encounter notable disparity in 3 cancers like KIRP (0.12), PRAD (0.07), and LUAD (0.11), emphasizing the need for optimization in leveraging RPPA data for consistent cancer molecular subtype classification. While certain modality-classifier combinations may achieve high macro ACC in classification tasks, this does not necessarily imply the absence of bias. For instance, classifiers trained on Array (GBM: 0.17), miRNA (UCEC: 0.16, LUSC: 0.1), and RNASeq (PCPG: 0.27) exhibit significant PR-RE disparity. These findings highlight that, despite strong overall performance in some cases, the presence of disparity must be carefully considered when using these modalities for cancer molecular subtype classification.

Certain modalities exhibit better alignment between macro PR and RE, achieving more consistent performance across cancers. RNASeq-based classifiers demonstrate low disparity in cancers such as ACC (0.09), LAML (0.04), and BLCA (0.01), showcasing their strength in maintaining balanced PR and RE for cancer classification. Similarly, classifiers trained on the CNV modality exhibit consistently low disparity across 19 cancers, although a slightly higher disparity is observed in GBM (0.13). This suggests that CNV-based classifiers are generally robust but may encounter challenges in achieving consistent classification for GBM. Moreover, miRNA-based classifiers show low disparity in diverse cancers such as PCPG, PRAD, COAD, and BRCA, demonstrating their robustness and stability. Additionally, classifiers trained on Exon modality maintain consistent performance with relatively low disparity in cancers such as COAD (0), and PRAD (0.02). These findings underline the potential of RNASeq, miRNA, Exon, and CNV data for achieving reliable and consistent molecular subtype classification across cancers. In summary, while modalities like RNASeq, miRNA, Exon, and CNV demonstrate consistent performance, challenges faced by classifiers using SNP, RPPA, and Array data highlight the need for further optimization and targeted strategies to improve their effectiveness in cancer molecular classification tasks.

In summary, classifiers trained on miRNA, RNASeq, Exon, Meth., and CNV data emerge as the most effective for cancer molecular subtype classification. miRNA-based classifiers achieve top performance in 5 cancers (BLCA, BRCA, KIRC, LAML, and LIHC), while CNV-based classifiers perform best in 4 cancers (KIRP, ESCA, SKCM, and KIRP). Classifiers utilizing RNASeq and Exon data also demonstrate strong performance, achieving top accuracy in 2 cancers each (RNASeq: PCPG, PRAD; Exon: COAD, THCA). These modalities consistently exhibit low disparity between precision and recall across various cancers, such as ACC, BLCA, and THCA, ensuring more balanced classification outcomes. In contrast, classifiers trained on SNP and RPPA data often show higher disparities and lower predictive performance, particularly in cancers such as ACC, LAML, and HNSC. While CNV-based classifiers are generally robust, a slightly higher disparity observed in GBM suggests potential challenges in maintaining consistent classification performance for this cancer. Similarly, classifiers using RPPA data encounter notable disparities in cancers like KIRP, LUSC, and LUAD, indicating the need for further optimization. These findings emphasize the importance of selecting data modalities based on their predictive capacity for specific cancers and addressing modality-specific biases to improve molecular subtype classification.

### 3.3 RQ IVa Which ML classifiers demonstrate reliable performance across all omics modalities?

In the previous subsection, we identified the omics data modalities that provide the most effective and accurate cancer molecular subtype classification by analyzing the maximum macro ACC and PR-RE disparity of ML and DL classifiers. However, it is equally important to assess how individual ML classifiers perform across each omics modality. To address this, we investigate RQ IVa by conducting an in-depth analysis of the median and inter-quartile range (IQR) of the macro ACC of 15 different ML classifiers across 8 distinct omics modalities for 20 cancers. In addition, based on median and IQR of macro ACC, classifiers are grouped into three categories: high-performing classifiers (high ACC and low IQR, indicating robustness and consistency), moderate-performing classifiers (high ACC but high IQR, variability despite strong performance), and low-performing classifiers (low ACC and high IQR, inconsistency and reduced reliability). These insights guide the selection of robust classifiers tailored to omics modalities for cancer molecular subtype classification.

Figure 4a illustrates median macro ACC and IQR of 15 distinct ML classifiers in terms of 8 different omics modalities across 20 different cancers, with detailed performance scores presented in Supplementary File 1 Table S3. On the basis of median macro ACC and IQR, 3 different performance groups are derived which are presented in Table 3. In the Array modality, XGB exhibits superior predictive performance, achieving a median macro ACC of 0.755 and an IQR of 0.085, indicating strong performance alongside low variability. SVM, GNB, and HGB also show competitive results, with median macro ACC values of 0.685, 0.675, and 0.670, respectively, though they exhibit higher variability (IQRs: 0.180, 0.215, and 0.195). LR follows closely with macro ACC of 0.670, with varying levels of consistency. Among the moderate performers, CNB and RF show a decline in performance, with macro ACC values of 0.592 and 0.590, and higher variability (IQRs: 0.328 and 0.347) showing inconsistencies across cancer subtypes. BNB and KNN perform even lower, with macro ACC values of 0.585 and 0.505, and high disparity in performance across different cases. The lowest-performing classifiers in the Array modality include CB, DT, and AB, with macro ACC values of 0.500, 0.490, and 0.455, respectively. QDA, LGBM, and GB show the lowest results, with median macro ACC values dropping as low as 0.245, 0.230, and 0.225, respectively, accompanied by low IQR values, which suggest instability in their predictions. These findings indicate that XGB, SVM, and GNB are among the most effective classifiers for the Array modality in cancer molecular subtype classification, while QDA, LGB, and GB struggle to provide reliable results.

**Figure 4:**
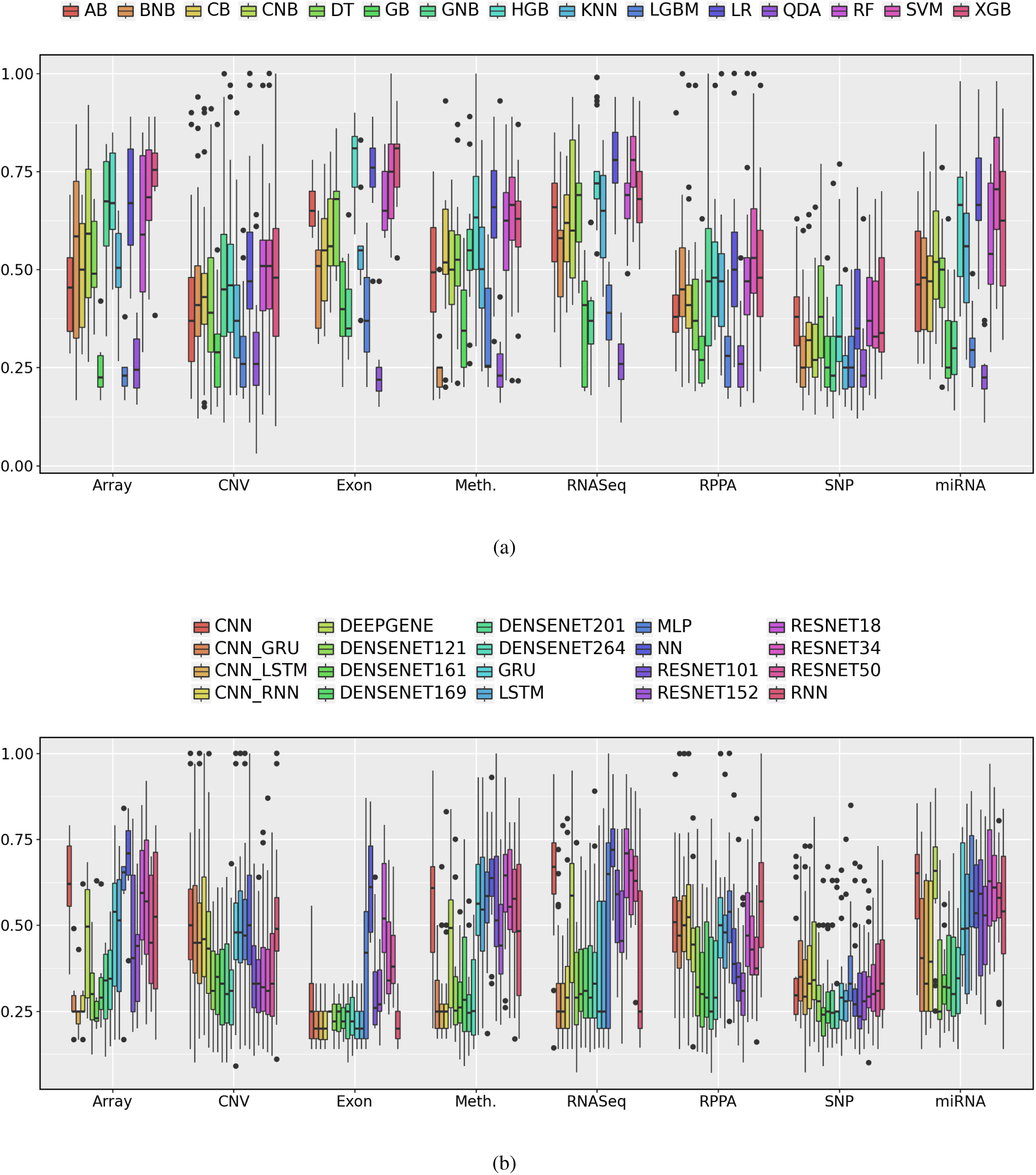
Modality-wise average macro ACC values for (a) ML (b) DL classifiers. For each omics modality, the average performance of a classifier is computed by aggregating its macro ACC values across all cancers. It highlights the comparative effectiveness of different classifiers w.r.t modalities for cancer molecular subtype classification.

**Table 3:** Three different categories of ML classifiers are assigned i.e., ☀ denotes classifiers with top performance for a modality, ● shows the moderate performing classifiers for a modality and × represents the lowest performance.

| Classifier | Array | CNV | Exon | Meth. | RNASeq | RPPA | SNP | miRNA |
| --- | --- | --- | --- | --- | --- | --- | --- | --- |
| **AB** | × | × | × | × | ● | × | × | × |
| **BNB** | ● | × | × | × | × | × | × | × |
| **CB** | × | × | ● | × | ● | × | × | × |
| **CNB** | ● | - | ● | × | ● | - | × | × |
| **DT** | × | × | × | ● | ★ | × | × | × |
| **GNB** | ★ | × | × | ● | × | × | × | × |
| **GB** | × | × | × | × | × | × | × | × |
| **HGB** | ★ | × | ★ | ★ | ★ | ● | × | ★ |
| **KNN** | ● | × | ● | × | ● | ● | × | ● |
| **LGBM** | × | × | × | × | × | × | × | × |
| **LR** | ★ | ● | ★ | ★ | ★ | ★ | × | ★ |
| **QDA** | × | × | × | × | × | × | × | × |
| **RF** | ● | ★ | × | ★ | ★ | ● | × | × |
| **SVM** | ★ | ★ | × | ★ | ★ | ★ | × | ★ |
| **XGB** | ★ | ● | ★ | × | ★ | ● | × | ● |

In the CNV modality, RF and SVM achieve the highest median macro ACC of 0.51 with IQRs of 0.180 and 0.175. XGB, and LR follow with macro ACC values of 0.48, and 0.47, and IQRs of 0.275, and 0.195. HGB and GNB perform moderately with macro ACC values of 0.46 and 0.45 and IQRs of 0.225 and 0.260. CB and BNB show slightly lower performance with median macro ACC values of 0.43 and 0.41 and IQRs of 0.130 and 0.180. DT, AdaBoost, and KNN perform worse with macro ACC values of 0.39, 0.37, and 0.37 and IQRs of 0.240, 0.215, and 0.185. GB, LGBM, and QDA show the lowest performance with macro ACC values of 0.29, 0.26, and 0.26 and moderate IQR values. RF, SVM, and XGB perform best in the CNV modality. QDA, LGBM, and GB show the lowest results. CNB is not applicable to CNV data due to negative values across all configurations of CNV modality.

For the SNP modality, none of the classifiers show reasonable performance, as all classifiers achieve low median macro ACC values with high variability. AB and DT perform the best but only achieve 0.38 macro ACC with inconsistent results (IQR: 0.125 and 0.235). RF, LR, XGB, and HGB fall within the 0.330.37 range, but all exhibit high variability. The remaining classifiers, including SVM, CB, CNB, BNB, GB, KNN, LBGM, QDA, and GNB, perform even worse, with macro ACC values dropping to 0.2300.330.

The RNASeq modality features LR and SVM as the top performers, both achieving a macro ACC of 0.78 and a moderate IQR of 0.130. HGB follows closely, with a macro ACC of 0.72 and a lower IQR of 0.070. DT and RF perform moderately, reaching a macro ACC of 0.69 and IQRs of 0.150 and 0.090, respectively. XGB achieves a 0.68 macro ACC with an IQR of 0.130, making it another competitive choice. AB, and KNN also show moderate performance, with macro ACC values between 0.66 and 0.65. CB and CNB perform moderately, with macro ACC values of 0.62 and 0.60 and IQRs of 0.170 and 0.352. The weakest classifiers, BNB, GB, LGBM, GNB, and QDA, achieve macro ACC values between 0.58 and 0.26, confirming their ineffectiveness for RNASeq-based cancer molecular subtype classification.

In the Exon modality, HGB and XGB achieve the highest performance, both with a macro ACC of 0.81 and IQRs of 0.13 and 0.11. LR follows closely with a macro ACC of 0.76 and a low IQR of 0.10. SVM performs well with a macro ACC of 0.75 but shows higher variability (IQR: 0.19). Among the moderate performers, DT, AB, and RF show macro ACC values of 0.68, 0.65, and 0.65, with IQRs between 0.09 and 0.19, demonstrating mixed levels of consistency. CNB, CB, and KNN achieve lower performance, with macro ACC values of 0.56 and 0.55, but KNN has the lowest IQR (0.06), indicating more stable predictions. The weakest classifiers include BNB, GB, LGBM, GNB, and QDA, all with macro ACC values below 0.51. QDA exhibits the worst performance, with a macro ACC of 0.22 and an IQR of 0.06, which shows its ineffectiveness for Exon-based cancer molecular subtype classification.

In the Meth. modality, LR and SVM perform the best, both achieving a macro ACC of 0.67 with moderate variability (IQR: 0.1725 and 0.1600). RF (0.62), HGB (0.633), and XGB (0.630) follow closely. GNB (0.55), and DT (0.525) perform moderately but with increased variability. CB, CNB, and KNN fall below 0.52, indicating weaker predictive capability. The lowest-performing classifiers include GB (0.34), LGBM (0.26), BNB (0.25), and QDA (0.23), confirming their ineffectiveness for Meth.-based classification.

In the RPPA modality, none of the classifiers perform well, with all showing low median macro ACC and high variability. SVM (0.53), and LR (0.50) achieve the highest scores but remain suboptimal. HGB, XGB, RF, and KNN perform similarly (0.480.47) with inconsistent results, especially GNB (IQR: 0.300). BNB, CatBoost, AdaBoost, DT, LGBM, GB, and QDA also fall below 0.46 which suggests the ineffectiveness for the use of ML classifiers for cancer molecular subtype classification.

In the miRNA modality, SVM (0.705) shows the best performance but with high variability (IQR: 0.235). HGB and LR (0.665) follow, with HGB exhibiting greater inconsistency (IQR: 0.25) than LR (0.16). XGB (0.625) perform moderately, though XGB shows high instability (IQR: 0.292). Lower performers include KNN (0.56), RF (0.54), CNB (0.52), and DT (0.50) while BNB, CB, and AB (0.480.46) demonstrate weaker predictive capacity. The worst classifiers, GNB, LightGBM, GBoost, and QDA (≤ 0.30), show their limited ability to handle miRNA modality for cancer molecular subtype classification.

Out of 16 different ML classifiers, only 6 i.e., LR, SVM, HGB, XGB, and RF consistently demonstrate high median macro ACC values with relatively low IQRs. Their strong predictive capability makes them the most suitable choices for cancer molecular subtype classification. In contrast, BNB, AB, CB, DT, GNB, GB, KNN, LGBM, MLP, and QDA demonstrate the lowest performance across all omics modalities. Their poor accuracy and high variability limit their effectiveness for cancer molecular subtype classification. These trends highlight the importance of balancing median macro ACC and IQR when selecting classifiers for specific modalities.

### 3.4 RQ IVb What DL classifiers demonstrate reliable performance across all omics modalities?

Following the insights of ML classifiers from the previous subsection, hereby we address RQ IVb to assess the performance of 20 different DL classifiers across 8 distinct omics modalities in terms of 20 different cancers based on independent testing. The grouping of classifiers into three categorieshigh-performing (high ACC and low IQR), moderate-performing (high ACC but high IQR), and low-performing (low ACC and high IQR)is done as explained in the previous subsection. This classification provides insights into the reliability of different DL models for cancer molecular subtype classification across omics modalities.

Figure 4b illustrates median macro ACC and IQR of 20 distinct DL classifiers across 8 different omics modalities in terms of 20 different cancers, with detailed performance scores presented in Supplementary File 1 Table S4. On the basis of median macro ACC and IQR, 3 different performance groups are derived which are presented in Table 4. In the Array modality, NN (0.71, IQR: 0.13) shows the best performance, followed by MLP (0.655, IQR: 0.067) and CNN (0.62, IQR: 0.175). ResNet18 (0.595) ResNet34 (0.57), and GRU (0.54) perform well but with high variability (IQR: 0.260.30). The lowest-performing classifiers include ResNet101, DenseNet models (264, 201, 121, 169), RNN, LSTM, DeepGene Transformer, Resnet50, CNN-GRU, CNN-RNN, CNN-LSTM, all with macro ACC ≤ 0.525, which proves their ineffectiveness for Array-based cancer molecular subtype classification.

**Table 4:** Three different categories of DL classifiers i.e., ☀ denotes classifiers with top performance for a modality, ● shows the moderate performing classifiers for a modality and × represents the lowest performance.

| Classifier | Array | CNV | Exon | Meth. | RNA | RPPA | SNP | miRNA |
| --- | --- | --- | --- | --- | --- | --- | --- | --- |
| **MLP** | • | • | × | • | • | ★ | × | ★ |
| **NN** | ★ | ★ | ★ | ★ | ★ | ★ | × | ★ |
| **CNN** | ★ | ★ | × | • | • | ★ | × | ★ |
| **ResNet18** | • | × | ★ | ★ | ★ | × | × | ★ |
| **ResNet34** | • | × | × | • | • | × | × | • |
| **GRU** | • | • | × | • | × | ★ | × | • |
| **RNN** | × | • | × | × | × | ★ | × | • |
| **LSTM** | × | • | × | × | × | • | × | • |
| **DEEPGENE** | × | × | × | × | × | • | × | ★ |
| **ResNet50** | × | × | • | • | • | × | × | × |
| **ResNet101** | × | × | × | × | × | × | × | × |
| **ResNet152** | × | × | × | × | × | × | × | × |
| **DenseNet101** | × | × | × | × | × | × | × | × |
| **DenseNet201** | × | × | × | × | × | × | × | × |
| **DenseNet121** | × | × | × | × | × | × | × | × |
| **DenseNet169** | × | × | × | × | × | × | × | × |
| **CNN-GRU** | × | × | × | × | × | • | × | × |
| **CNN-RNN** | × | × | × | × | × | ★ | × | • |
| **CNN-LSTM** | × | × | × | × | × | • | × | × |
| **DenseNet161** | × | × | × | × | × | × | × | × |

In the CNV modality, NN and CNN achieve the highest performance with a median macro ACC of 0.50, though CNN shows lower variability (IQR: 0.205) compared to NN (IQR: 0.260). GRU (0.48), RNN (0.49), LSTM (0.48), and MLP (0.47) fall into the moderate-performing category, with relatively moderate variability (IQR: 0.1550.210). The lowest-performing classifiers include CNN-RNN, CNN-GRU, CNN-LSTM, DeepGene transformer, ResNet (18, 34, 50, 101, 152), and DenseNet (121, 161, 169, 201, 264), all with macro ACC ≤ 0.46, which prove their limited reliability for CNV-based cancer molecular subtype classification.

In the Exon modality, NN achieves the highest performance with a macro ACC of 0.61, though its high IQR of 0.25 indicates some inconsistency. ResNet18 (0.52) and MLP (0.42) show moderate performance, but both have high IQR values of 0.26 and 0.37, which affects their stability. The lowest-performing classifiers include ResNet34, ResNet152, ResNet101, CNN, CNN-RNN, CNN-LSTM, CNN-GRU, DenseNet models (121, 161, 169, 201, 264), and recurrent models (GRU, LSTM, RNN). All have macro ACC values of 0.38 or lower, which confirms their poor performance in Exon-based classification.

In the Meth. modality, ResNet18 (0.65) and NN (0.64) achieve the highest performance, with low IQR values of 0.16 and 0.108, which makes them the most effective classifiers. CNN (0.609), MLP (0.585), RESNET50 (0.58), GRU (0.56), and RESNET34 (0.55) show moderate performance, but their high IQR values between 0.14 and 0.25 affect their stability. The lowest-performing classifiers include RNN, LSTM, ResNet101, ResNet152, DeepGene, DenseNet models (121, 161, 169, 201, 264), CNN-GRU, CNN-RNN, and CNN-LSTM. All have macro ACC values of 0.54 or lower, which confirms their poor performance in Meth.-based cancer molecular subtype classification

In the RNASeq modality, NN (0.72) and ResNet18 (0.71) achieve the highest median macro ACC, with IQR values of 0.1175 and 0.21, which makes them the most reliable classifiers. CNN (0.67), MLP (0.65), ResNet34 (0.66), and ResNet50 (0.63) show moderate performance, with IQR values of 0.15, 0.54, 0.19, and 0.13, which indicates some variability. The lowest-performing classifiers include ResNet101 (0.59), ResNet152 (0.454), DeepGene (0.586), DenseNet models (121: 0.29, 161: 0.30, 169: 0.31, 201: 0.29, 264: 0.33), LSTM (0.25), GRU (0.25), RNN (0.25), CNN-RNN (0.29), CNN-GRU (0.25), and CNN-LSTM (0.25). These classifiers have macro ACC values of 0.59 or lower, which confirms their poor performance in RNASeq-based classification.

In the RPPA modality, RNN (0.57), MLP (0.54), and CNN-RNN (0.52) achieve the highest median macro ACC, with IQR values ranging from 0.15-0.24. CNN-GRU (0.47), GRU (0.50), CNN (0.51), and CNN-LSTM (0.50) also perform well, with IQR values between 0.15 and 0.22. LSTM (0.479), ResNet18 (0.47), DeepGene (0.45), and ResNet34 (0.43) show moderate performance, but higher IQR values in some cases affect their consistency. The lowest-performing classifiers include DenseNet models (121: 0.32, 161: 0.30, 169: 0.29, 201: 0.25, 264: 0.29), ResNet models (50: 0.37, 101: 0.35, 152: 0.31), and NN (0.38). These classifiers have macro ACC values of 0.38 or lower which indicates their poor performance in RPPA-based cancer molecular subtype classification. In addition, none of the DL classifiers perform well with SNP. All DL classifiers achieve macro ACC values below 0.35, with high variability, confirming SNP as an ineffective modality for DL-based cancer molecular subtype classification classification. In the miRNA modality, ResNet18 (0.63), CNN (0.65), MLP (0.60), and DEEPGENE transformer (0.66) achieve the highest performance, with high variability (IQR: 0.14-0.27). CNN-RNN, RESNET34, RNN, GRU, and LSTM also perform well with median macro ACC 0.39-0.59, but CNN-RNN has the highest IQR (0.37). Moderate performers include DenseNet models (0.25-0.32), ResNet152 (0.53) and ResNet101 (0.59), with IQR values of 0.22 and 0.34 and low performers include CNN-GRU (0.41) and CNN-LSTM (0.33), with IQR values above 0.35.

DL classifiers show distinct performance trends across 8 omics modalities. NN and ResNet18 consistently achieve high median macro ACC with low IQR, making them the most reliable classifiers across multiple modalities, including

Array, CNV, Exon, Methylation, and RNASeq. CNN and MLP perform well in RNASeq, RPPA, Array, CNV, and miRNA, though with slightly higher variability. Hybrid models like CNN-RNN and CNN-GRU perform effectively in RPPA and miRNA, indicating their ability to handle complex omics structures. Moderate performers such as ResNet34, ResNet50, GRU, and LSTM exhibit reasonable median macro ACC but with higher variability, limiting their consistency. Low-performing classifiers, including DenseNet models, ResNet101, ResNet152, and recurrent models like RNN and LSTM, struggle across most modalities, showing poor accuracy and high IQR. SNP emerges as the least effective modality for DL classifiers, with all models failing to achieve reliable performance.

### 3.5 RQ V Which specific ML and DL classifiers provide consistent predictive performance w.r.t cancers?

In the previous subsections 3.3 and 3.4, we comprehensively discussed the performance trends of 15 ML and 20 DL classifiers across 8 distinct omics modalities based on independent testing. However, it is also important to analyze classifiers’ performance across different cancers to understand their generalizability and reliability in cancer molecular subtype classification. Since cancer subtypes exhibit distinct molecular characteristics, some classifiers may excel in specific cancers but struggle in others due to data complexity, sample heterogeneity, or imbalanced subtype distributions. To illustrate this, we investigate RQ V by selecting the highest predictive performance achieved by each classifier for each cancer, regardless of the omics modality based on independent testing. This analysis first identifies the top-performing ML and DL classifiers for each cancer type, followed by an evaluation of the most consistent classifiers that demonstrate robust performance across all cancers.

Figure 5a presents the highest macro ACC achieved by 15 ML classifiers across 20 cancers. SVM achieves the highest mean macro ACC of 0.78 and outperforms all other classifiers in 9 cancers: BLCA, BRCA, COAD, ESCA, KIRC, LIHC, SKCM, STAD, and UCEC. LR ranks second with a mean macro ACC of 0.77 and records the highest performance in 4 cancers: BLCA, ESCA, KIRP, and UCEC. XGB follows closely with a mean macro ACC of 0.75 and performs best in 3 cancers: GBM, LAML, and ESCA. Other boosting-based classifiers such as HGB and CB also achieve competitive results in some cancers i.e., PRAD, THCA, and ACC. These findings suggest that ensemble, SVM, LR, XGB, and HGB classifiers remain strong contenders for cancer molecular subtype classification across a diverse array of cancers.

**Figure 5:**
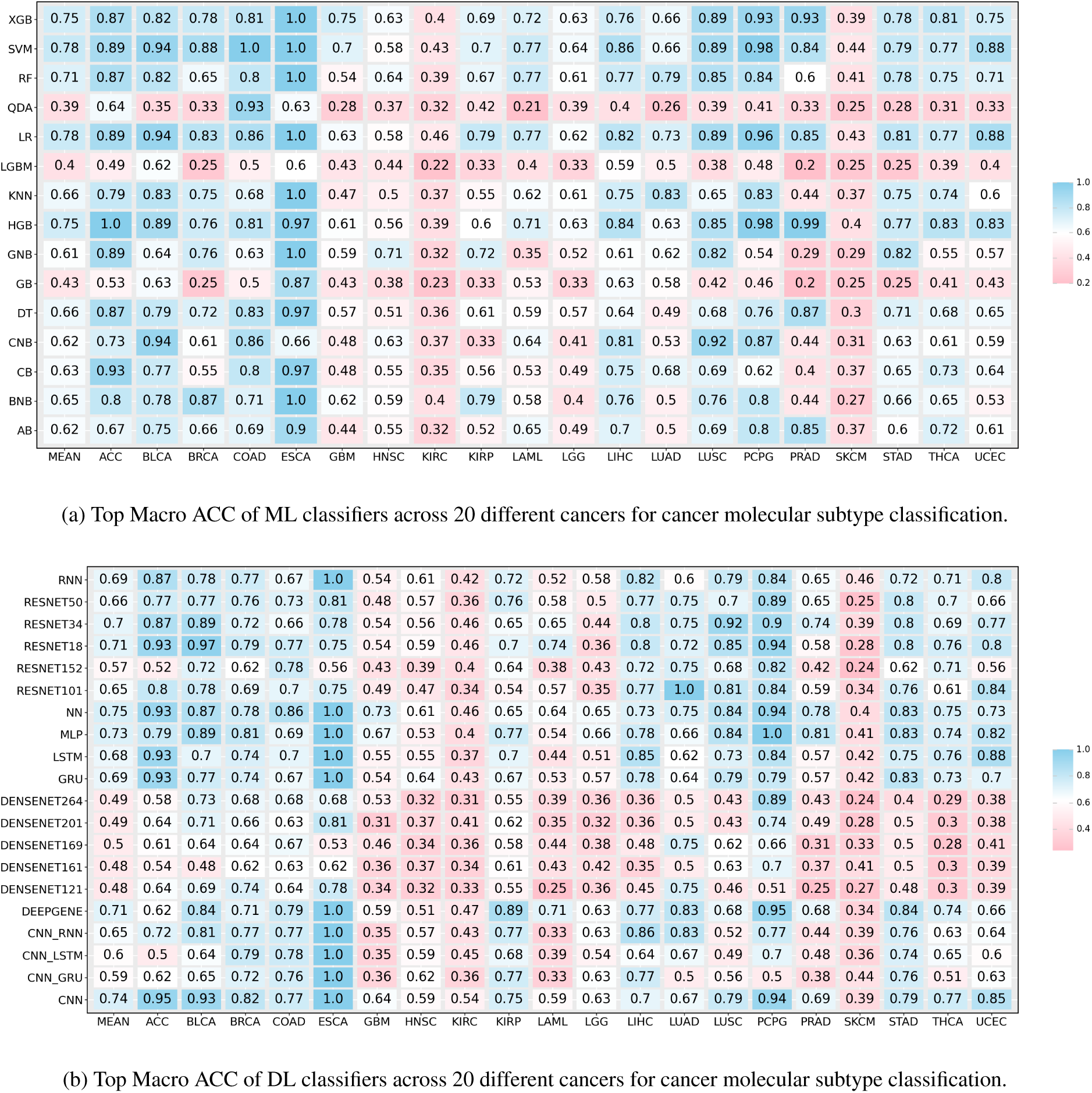
Across each cancer, the top performance of a classifier is shown regardless of the omics modality or dataset configuration to identify the most consistent classifiers for cancer molecular subtype classification.

Figure 5b presents the highest macro ACC achieved by 20 DL classifiers across 20 distinct cancers. NN achieves the highest mean macro ACC of 0.74, with the highest macro ACC in BLCA, GBM, LAML, and SKCM. MLP follows closely with an mean macro ACC of 0.72, having the highest macro ACC across 4 cancers namely, ESCA, LGG, PCPG, and PRAD. DeepGene transformer and CNN follow closely with a mean macro ACC of 0.71, with DeepGene transformer excelling in PCPG and KIRP, while CNN ranks the highest in ACC, BLCA, LUSC, and PCPG. Among ResNet architectures, ResNet18 achieves the best performance with a mean macro ACC of 0.70, excelling in BLCA, BRCA, and LUSC. GRU and LSTM achieve mean macro ACCs of 0.67 and 0.66, respectively, both performing best in BLCA and SKCM. DenseNet architectures (DenseNet264, DenseNet201, DenseNet169, DenseNet161, and DenseNet121) show moderate performance, with mean macro ACCs between 0.49 and 0.52, without dominating any particular cancer type. Hybrid models such as CNN-RNN, CNN-LSTM, and CNN-GRU perform moderately, with CNN-RNN reaching a mean macro ACC of 0.62, ranking highest for KIRP.

### 3.6 RQ VI Among top-performing classifiers in specific cancers, how do ML and DL methods compare in terms of consistency and suitability for cancer molecular subtype classification?

In the previous subsections, we provided insights into 17 different dataset configurations, where 5 configurations are identified as the most effective across 5/8 omics modalities. Then, in subsections 3.2, 3.3, 3.4, and 3.5, we evaluated 35 different ML and DL classifiers in terms of 20 distinct cancers and 8 different omics modalities, where 5 ML classifiers (SVM, LR, XGB, HGB, and RF) and 5 DL classifiers (NN, MLP, CNN, ResNet18, ResNet34, RNN, and DEEPGENE) are identified as the most reliable classifiers for cancer molecular subtype classification. Hereby, we address RQ VI by summarizing the top-performing combinations of ML and DL classifiers, dataset configurations, and omics modalities which provide a comprehensive foundation for optimal AI-driven cancer molecular subtype classification.

Table 5 shows the top-performing ML/DL classifiers across 20 cancers where DL classifiers outperform ML classifiers in 11 out of 20 cancers i.e., BLCA, ESCA, KIRP, KIRC, LGG, LUAD, LUSC, PCPG, SKCM, STAD, and UCEC. In contrast, ML models excel in 9 cancers, namely ACC, BRCA, COAD, GBM, HNSC, LAML, LIHC, PRAD, and TCHA. The performance differences between ML and DL classifiers can be attributed to key dataset characteristics such as the number of features, the number of samples, and the number of classes. These key attributes of the benchmark datasets are previously discussed in section 2.2.

**Table 5:**
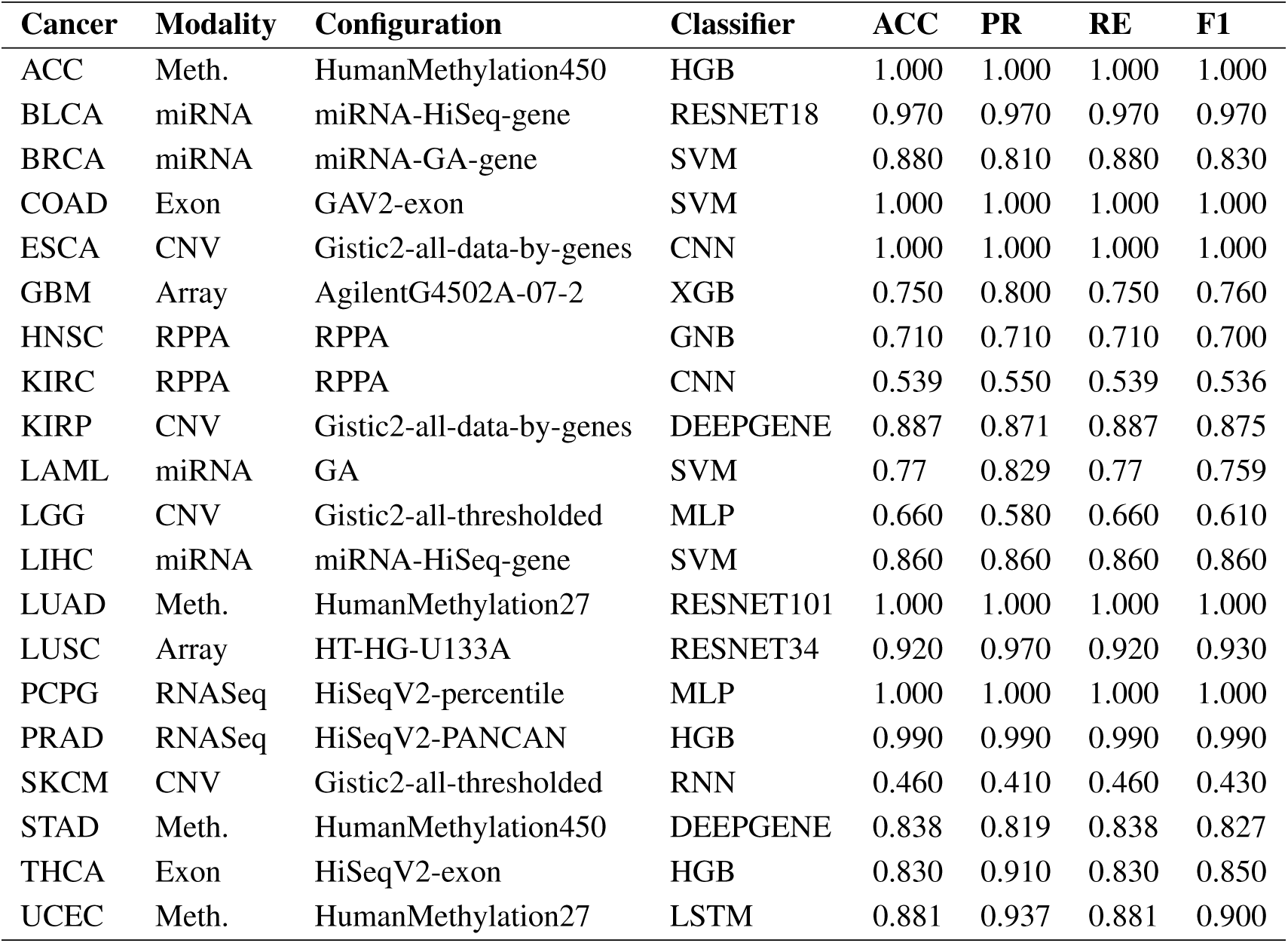
Top-Performing Classifier-Modality-Configuration Combinations for Cancer Molecular Subtype Classification.

ML models such as SVM, HGB, XGB, and RF tend to excel when datasets have fewer features, samples, and class labels. A lower feature count, typically between 131 and 10,000, reduces the need for deep feature extraction, making traditional ML models a more efficient choice. For instance, in BRCA (miRNA-GA-gene, SVM, ACC = 0.88) and UCEC (HumanMethylation27, LSTM, ACC = 0.88), where the feature count is relatively low (60015,000), SVM performs well by effectively separating classes with kernel methods without requiring complex hierarchical feature learning. ML models also outperform DL models when the datasets have a limited number of samples as DL classifiers require larger datasets to avoid overfitting. For instance, HNSC (RPPA, GNB, ACC = 0.71), and LIHC (miRNA, SVM, ACC = 0.86) where the total number of samples are 100-200, ML classifiers provides robust performance by effectively handling small-sample datasets.

DL models like ResNet18, ResNet34, ResNet101, CNN, RNN, and DEEPGENE excel with high-dimensional datasets (>20,000 features), larger sample sizes (≥ 150), and multi-class classification. They leverage hierarchical feature representations that traditional ML classifiers struggle to model. For example, in KIRP (CNV, DEEPGENE, ACC = 0.887) with 24,776 features and 161 samples, DL outperforms ML by capturing complex patterns despite the moderate sample size. Similarly, in LUAD (HumanMethylation27, ResNet101, ACC = 1.00) with 24,979 features and 226 samples, DL effectively handles high-dimensional data. LUSC (HT-HG-U133A, ResNet34, ACC = 0.92) further demonstrates DLs superiority in cancer molecular subtype classification (12,042 features, 104 samples). DL models generalize well with >150 samples where they prevent overfitting and capture intricate feature relationships. For instance, in PCPG (HiSeqV2-percentile, MLP, ACC = 1.00) with 20,501 features and 173 samples, and STAD (somatic mutation, DeepGene, ACC = 0.83) with 40,543 features and 383 samples, DL outperforms ML by modeling complex feature interactions across class distributions. Additionally, DL models perform well in datasets with six or more classes, effectively distinguishing cancer molecular subtypes. In LGG (7 classes, 513 samples), DL excels in multi-class classification, while in GBM (7 classes, 577 samples), ML (XGB) is an exception where sample size and feature dimensionality likely outweigh class complexity.

## 4 Discussion

This study presents a comprehensive benchmarking framework for evaluating the predictive performance of 35 different ML/DL classifiers across 153 datasets spanning 8 omics modalities and 20 cancer types for cancer molecular subtype classification. The results provide critical insights into the effectiveness of different omics modalities, data configurations, and AI classifiers for cancer molecular subtype classification. In this context, we discuss the key findings, their implications, and future directions for research in this domain.

The first critical step in designing effective and accurate predictive pipelines for cancer molecular subtype classification is the selection of optimal omics modalities and their configurations. Among data configurations, Gistic2-all-data-by-genes (CNV) and HiSeqV2 (RNASeq) consistently outperform others, emphasizing the value of granular, continuous genomic representations. HiSeqV2, HiSeqV2-PANCAN, and HiSeqV2-percentile show nearly identical performance, confirming RNASeqs robustness. In RPPA, unprocessed data outperforms RPPA-RBN by preserving essential signals. The array modality varies by configuration, while in Meth., HM27 surpasses HM450 due to a higher signal-to-noise ratio and lower overfitting risk. These findings highlight the impact of dataset configurations on classifier performance in cancer molecular subtype classification.

Our large-scale and comprehensive showed that miRNA, Meth., Exon, CNV, and RNASeq (gene expression), are the most informative modalities with the highest macro ACC in 14 cancers namely, BLCA, BRCA, LAML, LIHC, ESCA, KIRP, LGG, SKCM, COAD, THCA, PCPG, and PRAD. However, SNP and RPPA modalities showed lower predictive performance and higher PR-RE disparities which indicates challenges in leveraging these modalities for accurate cancer molecular subtype classification.

In terms of classifiers, large scale performance analyses identified SVM, LR, and XGB as the most consistent ML classifiers across multiple cancers, while ResNet18, CNN, and DEEPGENE emerged as the top-performing DL models. These classifiers demonstrated strong generalizability, achieving high macro ACC across diverse datasets and modalities. However, their performance varied depending on the cancer type and dataset characteristics, underscoring the need for modality-specific model selection. A comparison of ML and DL classifiers showed that ML models like SVM, LR, and XGB perform well in datasets with moderate feature counts and small sample sizes, effectively handling class imbalance and avoiding overfitting. In contrast, DL models such as ResNet, CNN, and DEEPGENE excel in high-dimensional datasets with larger sample sizes by capturing complex patterns through hierarchical feature representations. This distinction highlights the strengths of ML models for low-dimensional data and DL models for large-scale, multi-class classification tasks in omics research.

While DL models such as CNN, ResNet, and DEEPGENE have shown strong performance, Transformer-based models are emerging as a potential alternative for cancer molecular subtype classification i.e., DEEPGENE. Transformers have demonstrated state-of-the-art results in various domains but face significant challenges in omics applications. For instance, DEEPGENE relies on CNNs to transform features before self-attention is applied, which may lead to the loss of critical gene associations and other information. In addition, DEEPGENE is trained using supervised learning, requiring large labeled datasets. However, cancer subtype data is often limited, particularly for rare subtypes. Moreover, one major drawback of Transformers is their inability to handle extremely high-dimensional input directly. Standard architectures impose a maximum input length constraint (e.g., 512 tokens), which is insufficient for omics datasets containing tens of thousands of features (e.g., gene expression, methylation, CNV data). On the basis of such challenges, future work should investigate unsupervised or self-supervised pretraining approaches to reduce dependency on large labeled datasets, and effective embedding based strategies such that discriminatory and useful information is retained in the final representation of the data.

In terms of cancer molecular subtype classification, two different challenges are observed i.e., class imbalance, and modality bias. Class imbalance remains a significant challenge in cancer molecular subtype classification. For example, in BRCA, the Luminal A subtype (566 samples) is overrepresented compared to the Normal subtype (40 samples). This imbalance can lead to biased predictions, as classifiers may prioritize majority classes. Techniques such as stratified sampling and data augmentation should be explored to mitigate this issue. Modality-specific biases were also observed, particularly in SNP and RPPA datasets, where classifiers exhibited high PR-RE disparities. These biases may stem from noise, technical variability, or insufficient representation of certain subtypes in the data. Future work should focus on developing robust preprocessing pipelines and normalization techniques to address these challenges.

While this study provides valuable insights, several limitations should be acknowledged. First, the datasets used in this benchmark are primarily derived from TCGA, which may not fully capture the heterogeneity of cancer subtypes across different populations. Future studies should incorporate datasets from diverse sources, such as GEO and cBioPortal, to enhance generalizability. Second, the study focused on single-omics modalities, whereas multi-omics integration could provide a more comprehensive understanding of cancer molecular subtypes. Developing AI models capable of integrating multiple omics layers (e.g., combining CNV, methylation, and RNAseq) is a promising direction for future research.

Additionally, the study did not explore the impact of hyperparameter tuning on classifier performance. Optimizing hyperparameters for specific modalities and cancers could further improve predictive accuracy. Finally, the interpretability of AI models remains a critical challenge. While DL models often outperform ML classifiers, their “black-box” nature limits their adoption in clinical settings. Future work should focus on developing explainable AI (XAI) techniques to enhance the transparency and trustworthiness of these models.

## 5 Conclusion

This study in hand presents a comprehensive benchmarking analysis of AI-driven cancer molecular subtype classification across 20 different cancers using 8 distinct omics modalities and 35 ML/DL classifiers. Our findings highlight that certain omics modalities, such as RNASeq, miRNA, Exon, and CNV, provide more consistent and reliable performance in cancer molecular subtype classification, while others, such as SNP, RPPA, and Array, exhibit greater variability and require further optimization. Among classifiers, ML models like SVM, XGB, and HGB tend to perform well on smaller datasets with lower feature counts, whereas DL models such as ResNet18, CNN-RNN, and CNN-GRU demonstrate superior performance on high-dimensional datasets with large sample sizes. Additionally, the study underscores the disparity between precision and recall (PR-RE imbalance) as a key challenge in classification tasks, particularly for underperforming modalities. Overall, our benchmarking results provide critical insights into the strengths and limitations of different omics modalities and AI classifiers in cancer molecular subtype classification. By identifying the most effective classifier-modality combinations, this study serves as a foundation for developing robust and standardized AI pipelines for precision oncology. Future work should focus on mitigating modality-specific biases, improving performance on challenging data types, integrating multi-omics approaches to enhance classification accuracy and generalizability, and the application of transformer-based classifiers for cancer molecular subtype classification.

## 6 Availability of source code and requirements (optional, if code is present)

Source code can be provided at a reasonable request by the corresponding authors.

## 7 Data availability

The dataset(s) supporting the results of this article are available through UCSC Xena https://xenabrowser.net/datapages/.

## References

[1] Garima Mathur, Sumitra Nain, and Pramod Kumar Sharma. Cancer: an overview. Acad. J. Cancer Res, 8(1):01–09, 2015.

[2] Wolfgang Arthur Schulz. An introduction to human cancers. Molecular Biology of Human Cancers: An Advanced Student’s Textbook, pages 1–23, 2007.

[3] Jochen J Ott, Anke Ullrich, and Anthony B Miller. The importance of early symptom recognition in the context of early detection and cancer survival. European Journal of Cancer, 45(16):2743–2748, 2009.

[4] Third Edition. International classification of diseases for oncology. World Health Organization, 2020.

[5] Hyuna Sung, Jacques Ferlay, Rebecca L Siegel, Mathieu Laversanne, Isabelle Soerjomataram, Ahmedin Jemal, and Freddie Bray. Global cancer statistics 2020: Globocan estimates of incidence and mortality worldwide for 36 cancers in 185 countries. CA: a cancer journal for clinicians, 71(3):209–249, 2021.

[6] Henry HQ Heng, Joshua B Stevens, Steven W Bremer, Guo Liu, Batoul Y Abdallah, and J Ye Christine. Evolutionary mechanisms and diversity in cancer. Advances in cancer research, 112:217–253, 2011.

[7] Gloria H Heppner, William R Shapiro, and Joan K Rankin. Tumor heterogeneity. Pediatric Oncology 1: with a special section on Rare Primitive Neuroectodermal Tumors, pages 99–116, 1981.

[8] Li Ding, Matthew H Bailey, Eduard Porta-Pardo, Vesteinn Thorsson, Antonio Colaprico, Denis Bertrand, David L Gibbs, Amila Weerasinghe, Kuan-lin Huang, Collin Tokheim, et al. Perspective on oncogenic processes at the end of the beginning of cancer genomics. Cell, 173(2):305–320, 2018.

[9] Ju-Seog Lee. Exploring cancer genomic data from the cancer genome atlas project. BMB reports, 49(11):607, 2016.

[10] Jimmy Lin and Meng Li. Molecular profiling in the age of cancer genomics. Expert review of molecular diagnostics, 8(3):263–276, 2008.

[11] Miaolong Lu and Xianquan Zhan. The crucial role of multiomic approach in cancer research and clinically relevant outcomes. EPMA Journal, 9(1):77–102, 2018.

[12] Ahtisham Fazeel Abbasi, Muhammad Nabeel Asim, Sheraz Ahmed, Sebastian Vollmer, and Andreas Dengel. Survival prediction landscape: an in-depth systematic literature review on activities, methods, tools, diseases, and databases. Frontiers in Artificial Intelligence, 7:1428501, 2024.

[13] Marco Stricker, Muhammad Nabeel Asim, Andreas Dengel, and Sheraz Ahmed. Circnet: an encoder–decoder-based convolution neural network (cnn) for circular rna identification. Neural Computing and Applications, pages 1–12, 2022.

[14] Muhammad Nabeel Asim, Muhammad Ali Ibrahim, Ahtisham Fazeel, Andreas Dengel, and Sheraz Ahmed. Dnamp: a generalized dna modifications predictor for multiple species based on powerful sequence encoding method. Briefings in Bioinformatics, 24(1):bbac546, 2023.

[15] Muhammad Nabeel Asim, Muhammad Imran Malik, Christoph Zehe, Johan Trygg, Andreas Dengel, and Sheraz Ahmed. A robust and precise convnet for small non-coding rna classification (rpc-snrc). IEEE Access, 9:19379– 19390, 2020.

[16] Ahtisham Fazeel Abbasi, Muhammad Nabeel Asim, and Andreas Dengel. Transitioning from wet lab to artificial intelligence: a systematic review of ai predictors in crispr. Journal of Translational Medicine, 23:153, 2025.

[17] Jean-Emmanuel Bibault, Philippe Giraud, and Anita Burgun. Big data and machine learning in radiation oncology: state of the art and future prospects. Cancer letters, 382(1):110–117, 2016.

[18] Muhammad Nabeel Asim, Ahtisham Fazeel, Muhammad Ali Ibrahim, Andreas Dengel, and Sheraz Ahmed. Mp-vhppi: Meta predictor for viral host protein-protein interaction prediction in multiple hosts and viruses. Frontiers in Medicine, 9:1025887, 2022.

[19] Konstantina Kourou, Themis P Exarchos, Konstantinos P Exarchos, Michalis V Karamouzis, and Dimitrios I Fotiadis. Machine learning applications in cancer prognosis and prediction. Computational and structural biotechnology journal, 13:8–17, 2015.

[20] Muhammad Sufyan, Zeeshan Shokat, and Usman Ali Ashfaq. Artificial intelligence in cancer diagnosis and therapy: Current status and future perspective. Computers in Biology and Medicine, page 107356, 2023.

[21] Kritika Gaur and Miheer M Jagtap. Role of artificial intelligence and machine learning in prediction, diagnosis, and prognosis of cancer. Cureus, 14(11), 2022.

[22] Yanjiao Ren, Yimeng Gao, Wei Du, Weibo Qiao, Wei Li, Qianqian Yang, Yanchun Liang, and Gaoyang Li. Classifying breast cancer using multi-view graph neural network based on multi-omics data. Frontiers in Genetics, 15:1363896, 2024.

[23] Joung Min Choi and Heejoon Chae. mobrca-net: a breast cancer subtype classification framework based on multi-omics attention neural networks. BMC bioinformatics, 24(1):169, 2023.

[24] Asmaa M Hassan, Safaa M Naeem, Mohamed AA Eldosoky, and Mai S Mabrouk. Multi-omics-based machine learning for the subtype classification of breast cancer. Arabian Journal for Science and Engineering, pages 1–14, 2024.

[25] Lin Lin and Yongxia Bao. Development and validation of machine learning models for diagnosis and prognosis of lung adenocarcinoma, and immune infiltration analysis. Scientific Reports, 14(1):22081, 2024.

[26] Umesh Prasad, Soumitro Chakravarty, and Gyaneshwar Mahto. Lung cancer detection and classification using deep neural network based on hybrid metaheuristic algorithm. Soft Computing, 28(15):8579–8602, 2024.

[27] Anwar Khan and Boreom Lee. Deepgene transformer: Transformer for the gene expression-based classification of cancer subtypes. Expert Systems with Applications, 226:120047, 2023.

[28] Jordan Anaya, John-William Sidhom, Faisal Mahmood, and Alexander S Baras. Multiple-instance learning of somatic mutations for the classification of tumour type and the prediction of microsatellite status. Nature biomedical engineering, 8(1):57–67, 2024.

[29] Alexander Sarachakov, Andrey Tyshevich, Anna Belozerova, Ekaterina Postovalova, Alexander Bagaev, Vladimir Kushnarev, and Nathan Hale Fowler. An unsupervised h&e-based machine-learning approach for precise prediction of tumor microenvironment subtypes., 2024.

[30] Can Liu, Yuchen Duan, Qingqing Zhou, Yongkang Wang, Yong Gao, Hongxing Kan, and Jili Hu. A classification method of gastric cancer subtype based on residual graph convolution network. Frontiers in Genetics, 13:1090394, 2023.

[31] K Dhana Shree and S Logeswari. Odrnn: optimized deep recurrent neural networks for automatic detection of leukaemia. Signal, Image and Video Processing, 18(5):4157–4173, 2024.

[32] Youpeng Yang, Qiuhong Zeng, Gaotong Liu, Shiyao Zheng, Tianyang Luo, Yibin Guo, Jia Tang, and Yi Huang. Hierarchical classification-based pan-cancer methylation analysis to classify primary cancer. BMC bioinformatics, 24(1):465, 2023.

[33] Zarif L Azher, Anish Suvarna, Ji-Qing Chen, Ze Zhang, Brock C Christensen, Lucas A Salas, Louis J Vaickus, and Joshua J Levy. Assessment of emerging pretraining strategies in interpretable multimodal deep learning for cancer prognostication. BioData Mining, 16(1):23, 2023.

[34] M Shalini and S Radhika. Ig-ango: a novel ensemble learning algorithm for breast cancer prediction using genomic data. Evolving Systems, pages 1–20, 2024.

[35] Tanya Barrett, Stephen E Wilhite, Pierre Ledoux, Carlos Evangelista, Irene F Kim, Maxim Tomashevsky, Kimberly A Marshall, Katherine H Phillippy, Patti M Sherman, Michelle Holko, et al. Ncbi geo: archive for functional genomics data setsupdate. Nucleic acids research, 41(D1):D991–D995, 2012.

[36] Joung Min Choi, Chaelin Park, and Heejoon Chae. meth-semicancer: a cancer subtype classification framework via semi-supervised learning utilizing dna methylation profiles. BMC bioinformatics, 24(1):168, 2023.

[37] Xichao Wang, Hao Sun, Yongfei Dong, Jie Huang, Lu Bai, Zaixiang Tang, Songbai Liu, and Suning Chen. Development and validation of a cuproptosis-related prognostic model for acute myeloid leukemia patients using machine learning with stacking. Scientific Reports, 14(1):2802, 2024.

[38] Shimei Qin, Shibin Sun, Yahui Wang, Chao Li, Lei Fu, Ming Wu, Jinxing Yan, Wan Li, Junjie Lv, and Lina Chen. Immune, metabolic landscapes of prognostic signatures for lung adenocarcinoma based on a novel deep learning framework. Scientific Reports, 14(1):527, 2024.

[39] Abhibhav Sharma, Julia Debik, Bjørn Naume, Hege Oma Ohnstad, Tone F Bathen, and Guro F Giskeødegård. Comprehensive multi-omics analysis of breast cancer reveals distinct long-term prognostic subtypes. Oncogenesis, 13(1):22, 2024.

[40] Veronica Zelli, Andrea Manno, Chiara Compagnoni, Rasheed Oyewole Ibraheem, Francesca Zazzeroni, Edoardo Alesse, Fabrizio Rossi, Claudio Arbib, and Alessandra Tessitore. Classification of tumor types using xgboost machine learning model: a vector space transformation of genomic alterations. Journal of Translational Medicine, 21(1):836, 2023.

[41] Liancheng Jiang, Liye Jia, Yizhen Wang, Yongfei Wu, and Junhong Yue. Adap-bdcm: Adaptive bilinear dynamic cascade model for classification tasks on cnv datasets. Interdisciplinary Sciences: Computational Life Sciences, pages 1–19, 2024.

[42] Ankita Pandey and Arun Kumar. An integrated approach for breast cancer classification. Multimedia Tools and Applications, 82(21):33357–33377, 2023.

[43] Ş engül DOĞ AN, Burak TAŞ CI, and Türker TUNCER. Detection of acute lymphocytic leukemia (all) with a pre-trained deep learning model. 2023.

[44] Mahendran Botlagunta, Madhavi Devi Botlagunta, Madhu Bala Myneni, Deepa Lakshmi, Anand Nayyar, Jaithra Sai Gullapalli, and Mohd Asif Shah. Classification and diagnostic prediction of breast cancer metastasis on clinical data using machine learning algorithms. Scientific Reports, 13(1):485, 2023.

[45] Monika Lamba, Geetika Munjal, and Yogita Gigras. Ibcbml: interpreting breast cancer biomarker using machine learning. Health and Technology, pages 1–22, 2024.

[46] Lan Tian, Jiabao Wu, Wanting Song, Qinghuai Hong, Di Liu, Fei Ye, Feng Gao, Yue Hu, Meijuan Wu, Yi Lan, et al. Precise and automated lung cancer cell classification using deep neural network with multiscale features and model distillation. Scientific Reports, 14(1):10471, 2024.

[47] Sarfaraz Ahmed Mohammed, Senuka D Abeysinghe, and Anca L Ralescu. Feature selection and comparative analysis of breast cancer prediction using clinical data and histopathological whole slide images. Adv. Artif. Intell. Mach. Learn., 3(3):1494–1525, 2023.

[48] Jabed Omar Bappi, Mohammad Abu Tareq Rony, Mohammad Shariful Islam, Samah Alshathri, and Walid El-Shafai. A novel deep learning approach for accurate cancer type and subtype identification. IEEE Access, 2024.

[49] Wei Liu, Wei Wang, Hanyi Zhang, Miaoran Guo, Yingxin Xu, and Xiaoqi Liu. Development and validation of multi-omics thymoma risk classification model based on transfer learning. Journal of Digital Imaging, 36(5):2015–2024, 2023.

[50] Upeka Vianthi Somaratne, Kok Wai Wong, Jeremy Parry, and Hamid Laga. The use of generative adversarial networks for multi-site one-class follicular lymphoma classification. Neural Computing and Applications, 35(28):20569–20579, 2023.

[51] J. Ross Quinlan. Induction of decision trees. Machine learning, 1:81–106, 1986.

[52] Yali Amit and Donald Geman. Shape quantization and recognition with randomized trees. Neural computation, 9(7):1545–1588, 1997.

[53] Leo Breiman. Random forests. Machine learning, 45:5–32, 2001.

[54] Jerome H Friedman. Greedy function approximation: a gradient boosting machine. Annals of statistics, pages 1189–1232, 2001.

[55] Aleksei Guryanov. Histogram-based algorithm for building gradient boosting ensembles of piecewise linear decision trees. In Analysis of Images, Social Networks and Texts: 8th International Conference, AIST 2019, Kazan, Russia, July 17–19, 2019*, Revised Selected Papers 8*, pages 39–50. Springer, 2019.

[56] Yoav Freund and Robert E Schapire. A decision-theoretic generalization of on-line learning and an application to boosting. Journal of computer and system sciences, 55(1):119–139, 1997.

[57] Guolin Ke, Qi Meng, Thomas Finley, Taifeng Wang, Wei Chen, Weidong Ma, Qiwei Ye, and Tie-Yan Liu. Lightgbm: A highly efficient gradient boosting decision tree. Advances in neural information processing systems, 30, 2017.

[58] Liudmila Prokhorenkova, Gleb Gusev, Aleksandr Vorobev, Anna Veronika Dorogush, and Andrey Gulin. Catboost: unbiased boosting with categorical features. Advances in neural information processing systems, 31, 2018.

[59] Leif E Peterson. K-nearest neighbor. Scholarpedia, 4(2):1883, 2009.

[60] Thorsten Joachims. Svmlight: Support vector machine. SVM-Light Support Vector Machine http://svmlight.joachims.org/*, University of Dortmund*, 19(4):25, 1999.

[61] Geoffrey J McLachlan. Discriminant analysis and statistical pattern recognition. John Wiley & Sons, 2005.

[62] Jerome Cornfield, Tavia Gordon, and Willie W Smith. Quantal response curves for experimentally uncontrolled variables. Bull Int Stat Inst, 38(3):97–115, 1961.

[63] S Walter and D Duncan. Estimation of the probability of an event as a function of several variables. Biometrika, 54(1-2):167–179, 1967.

[64] Manjunath Jogin, MS Madhulika, GD Divya, RK Meghana, S Apoorva, et al. Feature extraction using convolution neural networks (cnn) and deep learning. In 2018 3rd IEEE international conference on recent trends in electronics, information & communication technology (RTEICT), pages 2319–2323. IEEE, 2018.

[65] Jiuxiang Gu, Zhenhua Wang, Jason Kuen, Lianyang Ma, Amir Shahroudy, Bing Shuai, Ting Liu, Xingxing Wang, Gang Wang, Jianfei Cai, et al. Recent advances in convolutional neural networks. Pattern recognition, 77:354–377, 2018.

[66] Larry R Medsker, Lakhmi Jain, et al. Recurrent neural networks. Design and Applications, 5(64-67):2, 2001.

[67] Marina Sokolova, Nathalie Japkowicz, and Stan Szpakowicz. Beyond accuracy, f-score and roc: a family of discriminant measures for performance evaluation. In Australasian joint conference on artificial intelligence, pages 1015–1021. Springer, 2006.

[68] Mario Deng, Johannes Brägelmann, Joachim L Schultze, and Sven Perner. Web-tcga: an online platform for integrated analysis of molecular cancer data sets. BMC bioinformatics, 17:1–7, 2016.

[69] Katarzyna Tomczak, Patrycja Czerwińska, and Maciej Wiznerowicz. Review the cancer genome atlas (tcga): an immeasurable source of knowledge. Contemporary Oncology/Współczesna Onkologia, 2015(1):68–77, 2015.

[70] John N Weinstein, Eric A Collisson, Gordon B Mills, Kenna R Shaw, Brad A Ozenberger, Kyle Ellrott, Ilya Shmulevich, Chris Sander, and Joshua M Stuart. The cancer genome atlas pan-cancer analysis project. Nature genetics, 45(10):1113–1120, 2013.

[71] Antonio Colaprico, Tiago C Silva, Catharina Olsen, Luciano Garofano, Claudia Cava, Davide Garolini, Thais S Sabedot, Tathiane M Malta, Stefano M Pagnotta, Isabella Castiglioni, et al. Tcgabiolinks: an r/bioconductor package for integrative analysis of tcga data. Nucleic acids research, 44(8):e71–e71, 2016.

[72] Kevin P Murphy, et al. Naive bayes classifiers. University of British Columbia, 18(60):1–8, 2006.

[73] Andrew McCallum, Kamal Nigam, et al. A comparison of event models for naive bayes text classification. In AAAI-98 workshop on learning for text categorization, volume 752, pages 41–48. Madison, WI, 1998.

[74] Xin Shao, Ning Lv, Jie Liao, Jinbo Long, Rui Xue, Ni Ai, Donghang Xu, and Xiaohui Fan. Copy number variation is highly correlated with differential gene expression: a pan-cancer study. BMC medical genetics, 20:1–14, 2019.

[75] Craig H Mermel, Steven E Schumacher, Barbara Hill, Matthew L Meyerson, Rameen Beroukhim, and Gad Getz. Gistic2. 0 facilitates sensitive and confident localization of the targets of focal somatic copy-number alteration in human cancers. Genome biology, 12:1–14, 2011.

[76] Yuan Chun Ding, Hanbing Song, Aaron W Adamson, Daniel Schmolze, Donglei Hu, Scott Huntsman, Linda Steele, Carmina S Patrick, Shu Tao, Natalie Hernandez, et al. Profiling the somatic mutational landscape of breast tumors from hispanic/latina women reveals conserved and unique characteristics. Cancer research, 83(15):2600– 2613, 2023.

[77] Marie-Agnès Dillies, Andrea Rau, Julie Aubert, Christelle Hennequet-Antier, Marine Jeanmougin, Nicolas Servant, Céline Keime, Guillemette Marot, David Castel, Jordi Estelle, et al. A comprehensive evaluation of normalization methods for illumina high-throughput rna sequencing data analysis. Briefings in bioinformatics, 14(6):671–683, 2013.

[78] Günter P Wagner, Koryu Kin, and Vincent J Lynch. Measurement of mrna abundance using rna-seq data: Rpkm measure is inconsistent among samples. Theory in biosciences, 131:281–285, 2012.

[79] Malachi Griffith, Christopher A Miller, Obi L Griffith, Kilannin Krysiak, Zachary L Skidmore, Avinash Ramu, Jason R Walker, Ha X Dang, Lee Trani, David E Larson, et al. Optimizing cancer genome sequencing and analysis. Cell systems, 1(3):210–223, 2015.

[80] Minju Ha and V Narry Kim. Regulation of microrna biogenesis. Nature reviews Molecular cell biology, 15(8):509–524, 2014.

[81] Haley J Abel, David E Larson, Colby Chiang, Indraniel Das, Krishna L Kanchi, Ryan M Layer, Benjamin M Neale, William J Salerno, Catherine Reeves, Steven Buyske, et al. Mapping and characterization of structural variation in 17,795 deeply sequenced human genomes. *bioRxiv*, page 508515, 2018.

[82] Kyle Ellrott, Matthew H Bailey, Gordon Saksena, Kyle R Covington, Cyriac Kandoth, Chip Stewart, Julian Hess, Singer Ma, Kami E Chiotti, Michael McLellan, et al. Scalable open science approach for mutation calling of tumor exomes using multiple genomic pipelines. Cell systems, 6(3):271–281, 2018.

[83] Cristian Coarfa, Sandra L Grimm, Kimal Rajapakshe, Dimuthu Perera, Hsin-Yi Lu, Xuan Wang, Kurt R Christensen, Qianxing Mo, Dean P Edwards, and Shixia Huang. Reverse-phase protein array: Technology, application, data processing, and integration. Journal of biomolecular techniques: JBT, 32(1):15, 2021.

[84] Jun Li, Yiling Lu, Rehan Akbani, Zhenlin Ju, Paul L Roebuck, Wenbin Liu, Ji-Yeon Yang, Bradley M Broom, Roeland GW Verhaak, David W Kane, et al. Tcpa: a resource for cancer functional proteomics data. Nature methods, 10(11):1046–1047, 2013.

[85] Zhenxing Wang, XiaoLiang Wu, and Yadong Wang. A framework for analyzing dna methylation data from illumina infinium humanmethylation450 beadchip. BMC bioinformatics, 19:15–22, 2018.

[86] Christina Thirlwell, Marianne Eymard, Andrew Feber, Andrew Teschendorff, Kerra Pearce, Matthias Lechner, Martin Widschwendter, and Stephan Beck. Genome-wide dna methylation analysis of archival formalin-fixed paraffin-embedded tissue using the illumina infinium humanmethylation27 beadchip. Methods, 52(3):248–254, 2010.

87. K Tomczak, Czerwinska p, wiznerowicz m (2015) the cancer genome atlas (tcga): an immeasurable source of knowledge. Contemp Oncol, 19:A68.

[88] Shujun Huang, Nianguang Cai, Pedro Penzuti Pacheco, Shavira Narrandes, Yang Wang, and Wayne Xu. Applications of support vector machine (svm) learning in cancer genomics. Cancer genomics & proteomics, 15(1):41–51, 2018.

[89] Martin Riedmiller. Advanced supervised learning in multi-layer perceptronsfrom backpropagation to adaptive learning algorithms. Computer Standards & Interfaces, 16(3):265–278, 1994.

[90] Jianxin Wu. Introduction to convolutional neural networks. National Key Lab for Novel Software Technology. Nanjing University. China, 5(23):495, 2017.

[91] Gao Huang, Zhuang Liu, Laurens Van Der Maaten, and Kilian Q Weinberger. Densely connected convolutional networks. In Proceedings of the IEEE conference on computer vision and pattern recognition, pages 4700–4708, 2017.

[92] Kaiming He, Xiangyu Zhang, Shaoqing Ren, and Jian Sun. Identity mappings in deep residual networks. In Computer Vision–ECCV 2016: 14th European Conference, Amsterdam, The Netherlands, October 11–14, 2016*, Proceedings, Part IV 14*, pages 630–645. Springer, 2016.

[93] Alex Graves and Alex Graves. Long short-term memory. Supervised sequence labelling with recurrent neural networks, pages 37–45, 2012.

[94] KE ArunKumar, Dinesh V Kalaga, Ch Mohan Sai Kumar, Masahiro Kawaji, and Timothy M Brenza. Forecasting of covid-19 using deep layer recurrent neural networks (rnns) with gated recurrent units (grus) and long shortterm memory (lstm) cells. Chaos, Solitons & Fractals, 146:110861, 2021.

[95] Jerome Friedman. The elements of statistical learning: Data mining, inference, and prediction. *(No Title)*, 2009.

[96] Željko Ivezić, Andrew J Connolly, Jacob T VanderPlas, and Alexander Gray. Statistics, data mining, and machine learning in astronomy: a practical Python guide for the analysis of survey data, volume 8. Princeton University Press, 2020.

[97] Marina Sokolova and Guy Lapalme. A systematic analysis of performance measures for classification tasks. Information processing & management, 45(4):427–437, 2009.

[98] Konstantinos Sechidis, Grigorios Tsoumakas, and Ioannis Vlahavas. On the stratification of multi-label data. In Machine Learning and Knowledge Discovery in Databases: European Conference, ECML PKDD 2011, Athens, Greece, September 5-9, 2011, Proceedings, Part III 22, pages 145–158. Springer, 2011.

[99] Tzu-Tsung Wong. Performance evaluation of classification algorithms by k-fold and leave-one-out cross validation. Pattern recognition, 48(9):2839–2846, 2015.

